# Epigenetic Control of Hundreds of Chromosome-Associated lncRNA Genes Essential for Replication and Stability

**DOI:** 10.1101/2022.04.25.489405

**Authors:** Michael B. Heskett, Athanasios E. Vouzas, Leslie G. Smith, Phillip A. Yates, Christopher Boniface, Eric E. Bouhassira, Paul Spellman, David M. Gilbert, Mathew J. Thayer

## Abstract

ASARs are long noncoding RNA genes that control replication timing of entire human chromosomes in *cis*. The three known ASAR genes are located on human chromosomes 6 and 15, and are essential for chromosome integrity. To identify ASARs on all human chromosomes we utilized a set of distinctive ASAR characteristics that allowed for the identification of hundreds of autosomal loci with epigenetically controlled, allele-restricted behavior in expression and replication timing of coding and noncoding genes, and is distinct from genomic imprinting. Disruption of noncoding RNA genes at five of five tested loci resulted in chromosome-wide delayed replication and chromosomal instability, validating their ASAR activity. In addition to the three known essential *cis*-acting chromosomal loci, origins, centromeres, and telomeres, we propose that all mammalian chromosomes also contain “Inactivation/Stability Centers” that display allele-restricted epigenetic regulation of protein coding and noncoding ASAR genes that are essential for replication and stability of each chromosome.

## Introduction

For the vast majority of mammalian DNA, homologous regions on chromosome pairs replicate in a highly synchronized manner ^1-3^. However, genetic disruption of non-protein coding ASAR (“<u>AS</u>ynchronous replication and <u>A</u>utosomal <u>R</u>NA”) genes causes a delay in replication timing on individual human chromosomes in *cis*, resulting in highly asynchronous replication between pairs of autosomes ^4-6^. The first ASAR genes were identified from a genetic screen designed to identify loci on human chromosomes that when disrupted resulted in chromosome-wide delayed replication ^4,6,7^. This screen identified five balanced translocations, affecting eight different autosomes, all displaying delayed replication along the length of the chromosomes ^7^. Characterization of two of the translocation breakpoints identified discrete *cis*-acting loci where translocations or deletions resulted in delayed replication ^4,6^. Molecular examination of the disrupted loci identified two lncRNA genes located on chromosomes 6 and 15, and were named *ASAR6* and *ASAR15*, respectively ^4,6^. The ASAR6 and ASAR15 lncRNAs are extremely long (>200 kb), and remain associated with the chromosome territories where they are transcribed ^4,6,8-10^. These studies defined the first *cis*-acting loci that control replication timing and structural stability of individual human autosomes ^4-6^.

One unusual characteristic of ASAR genes is that they are expressed from only one allele ^4-6^. In contrast, the majority of genes on mammalian autosomes are expressed from both alleles, *i. e*. they display bi-allelic or balance expression. <u>A</u>llelic <u>E</u>xpression Imbalance (AEI) of protein coding and noncoding genes is well established, and can arise from several distinct mechanisms. For example, AEI can arise due to heterozygosity at DNA sequence polymorphisms within *cis*-acting elements that influence the efficiency with which a gene will be transcribed [*i. e.* <u>e</u>xpression <u>Q</u>uantitative <u>T</u>rait <u>L</u>oci or eQTL; reviewed in ^11,12^]. AEI may also occur through epigenetic programming to regulate gene dosage or to provide “exquisite specificity”, where the most extreme form of AEI is referred to as mono-allelic expression [reviewed in ^13-17^]. Genomic imprinting is a well-established form of epigenetically programmed AEI occurring in a parent of origin specific manner [reviewed in ^18,19^]. Alternatively, epigenetically programmed AEI occurring in a random manner with respect to parent of origin has been observed for as many as 10% of autosomal genes [e. g. olfactory receptors, immunoglobulins, and T cell receptors; ^20-23^]. The three known ASAR genes display random epigenetically programed AEI ^4-6,9^.

Another unusual characteristic of ASAR genes is that they display <u>AS</u>ynchronous <u>R</u>eplication <u>T</u>iming (ASRT) between alleles ^4-6,9,24^. In contrast, the timing of DNA replication on autosome pairs occurs in a highly synchronous manner ^1,2^. ASRT can arise by different mechanisms. For example, ASRT can be caused by heterozygosity at DNA sequence polymorphisms that dictate the time during S phase that a locus will be replicated [*i. e*. <u>r</u>eplication timing <u>Q</u>uantitative <u>T</u>rait <u>L</u>oci or rtQTL; ^25-27^]. In addition, ASRT occurs at regions of the genome that display epigenetically programmed AEI. Thus, both imprinted and random AEI genes located on autosomes display ASRT ^1,13,28-32^. The three known ASAR genes display random epigenetically programed ASRT between alleles ^4-6,9,24^.

ASAR RNAs share similarities with the vlinc (<u>v</u>ery long intergenic <u>n</u>on-<u>c</u>oding) class of RNAs. The vlincRNAs were characterized as RNA Polymerase II products that are nuclear, non-spliced, non-polyadenylated transcripts of >50 kb of contiguously expressed sequence that are not associated with protein coding genes ^33^. There are currently >2,700 annotated human vlincRNAs, which are expressed in a highly cell type-restricted manner ^33-36^. The annotated vlincRNAs have been referred to as genomic “Dark Matter” because the vast majority of them have unknown function. However, they occupy >10% of the human genome and represent >50% of the non-ribosomal RNA within the cell ^33-36^. Previously, we found that the genomic region annotated as expressing vlinc273 (>185 kb in length) has all of the physical characteristics that are shared between *ASAR6* and *ASAR15*, and CRISPR/Cas9-mediated deletion of the vlinc273 genomic locus resulted in delayed replication of chromosome 6, indicating that vlinc273 is an ASAR [which we named *ASAR6-141; ^5^*]. *ASAR6-141* resides within a cluster of 6 vlincRNA genes, with all 6 being expressed in a partially overlapping set of human tissues ^5,36^. Furthermore, the original *ASAR6* gene resides within a larger ∼1mb genomic region of ASRT that also contains two other vlincRNA genes, which are also expressed in different human tissues ^5,9^. Taken together, these observations raise the intriguing possibility that other vlincRNAs are also ASARs, and that multiple ASARs can be clustered at the same locus, expressed in different cell types and associated with the same ASRT region.

Deletion and ectopic integration analyses demonstrated that the chromosome-wide effects on replication timing of *ASAR6* and *ASAR15* map to the antisense strand of LINE1 (L1) retrotransposons located within the ASAR6 and ASAR15 RNAs ^10^. Targeting the L1 sequences, using oligonucleotide-directed RNA degradation, revealed a functional role for the L1 sequences within ASAR6 RNA in controlling chromosome-wide replication timing ^10^. Previous support for a role for L1s in epigenetically programmed AEI came from the observation that L1s are present at a relatively high local concentration (>18%) near both imprinted and random AEI genes located on autosomes ^37^. The three known ASAR genes contain >30% L1 sequences within the transcribed regions ^4-6^.

Altogether, our findings have suggested that ASARs may be ubiquitous essential *cis*-acting elements of chromosome replication and integrity. Indeed, we previously found that ∼2.5% of chromosome translocation products, induced by two different mechanisms (ionizing radiation or Cre/loxP), result in delayed replication timing of entire human chromosomes ^7,38^. Two of the Cre/LoxP-induced translocations with delayed replication were further characterized and found to have disrupted ASAR genes ^4,6^, suggesting that ∼2.5% of the human genome encodes ASARs. Here, we directly tested this hypothesis by taking advantage of several notable characteristics of the three known ASARs to identify additional ASAR genes on human autosomes: 1) long contiguously transcribed regions of >180 kb, 2) epigenetically regulated AEI; 3) epigenetically regulated ASRT; 4) high density of L1 sequences; and 5) spatial retention of the RNA on the parent chromosome. We used RNAseq, Repli-seq, and RNA-DNA FISH assays on single cell-derived clones from lymphoblastoid <u>c</u>ell lines (LCLs) isolated from two unrelated individuals, both with haplotype phased genomes, to identify additional ASAR candidates on human autosomes. We chose female cells for this analysis to take advantage of X chromosome inactivation as an internal control for epigenetically controlled AEI and ASRT that occurs between the active and inactive X chromosomes. We identified hundreds of autosomal loci that display epigenetically controlled AEI and ASRT that is comparable to that observed on the X chromosome. Genetic deletion assays validated the biological activity of five out of five tested ASAR candidates. Our work identifies a novel, widespread epigenetic program that occurs at hundreds of autosomal loci, effects AEI and ASRT of both noncoding and protein coding genes, has profound roles in chromosome structure stability, is predominant over the *cis*-effects of rtQTLs, and is distinct from genomic imprinting.

## Results

### Transcribed Loci with Allelic Expression Imbalance are widespread on autosomes

One complicating factor associated with both vlinc and ASAR RNAs is their extreme length. Thus, determining if these long RNA species represent single >50 kb contiguous transcripts or represent multiple overlapping transcripts with different start and stop sites is difficult using existing technologies. The vlincRNAs were annotated using tiling of contiguous RNAseq reads across a given locus allowing gaps in the contigs of ≤5 kb to accommodate repetitive elements, e. g. ALUs and LINEs, that don’t allow for unique mapping of the RNAseq reads to the genome ^34,36^. This complication is also evident for all of the known ASAR RNAs, where the long length (>180 kb) and high L1 content (>30%) make annotation of the transcripts difficult. Given these limitations we refer to regions of the genome with >50 kb of contiguous transcription of non-coding DNA as <u>T</u>ranscribed <u>L</u>oci (TL) throughout this manuscript.

We first sought to identify TL that are expressed in a human lymphoblastoid cell line (LCL). For this analysis we used publicly available data from a nuclear-enriched, ribosomal RNA depleted, strand-specific, RNA-seq dataset from GM12878 [see ENCODE; ^39^]. Using a strategy of merging contiguous reads, ^34,40^ a strand-specific call set of 1,570 TL was defined. An example of a TL is shown in Figure 1A, and shows the genomic location and the strand-specific, contiguous coverage of the RNAseq reads for TL:1-187. This TL covers ∼500 kb of genomic DNA, contains two RefSeq lincRNAs (LINC01036 and LINC01037), is not associated with any known protein coding gene, and contains 562 heterozygous SNPs within the transcribed region. Next, utilizing a fully haplotype-phased reference genotype (*i. e.* one haplotype block spanning each chromosome) for GM12878 to enumerate informative allele-specific reads ^41^ we assessed AEI of all 1,570 TL. GM12878 cells are female and represent a poly-clonal pool of EBV transformed lymphocytes. Random X chromosome inactivation among different cells in the population would be indicated by an equal number of RNA-seq reads from the maternal (haplotype 1) and paternal (haplotype 2) X chromosomes. However, deviation from equal inactivation of each parental allele is common in the general female population, and is known as skewing ^42,43^. We found dramatic AEI of both protein coding and non-coding RNAs expressed from the X chromosome, indicating that the LCL pool of cells in GM12878 has skewed X inactivation (see Table S1). We identified 78 TL that were expressed from the X chromosome, and 60 of these TL show AEI (Fig. 1B and 1C). The distribution of AEI of the X linked TL also revealed the presence of 18 TL with equivalent expression from both X chromosomes. These bi-allelic TL are located in regions of the X chromosome that are known to escape X inactivation [Table S1; and see ^43-45^]. Given the dramatic skew in X inactivation, and anticipating a similar skew in random AEI on autosomes, we next assessed the autosomal TL for allele-specific expression. In contrast to the X chromosome TL, the autosome-wide distribution of AEI is consistent with bi-allelic expression for the majority of TL, but 300 unique TL (encoded by ∼90 mb or ∼3% of the genome) have AEI that significantly deviated from the null distribution of equal expression from both alleles (Fig. 1B and 1C; and Table 1). These TL contain a high L1 content compared to protein coding genes (Fig. 1D), which is one of the characteristics of the known ASARs. The TL with AEI are distributed along the length of every autosome pair and are expressed in a reciprocal pattern (see Table S1). Figure 1E shows allelic expression analysis of all of the TL on chromosome 1, and highlights the 19 TL that display AEI, including TL:1-187 (see Fig. 1A and 1C).

**Figure 1.**
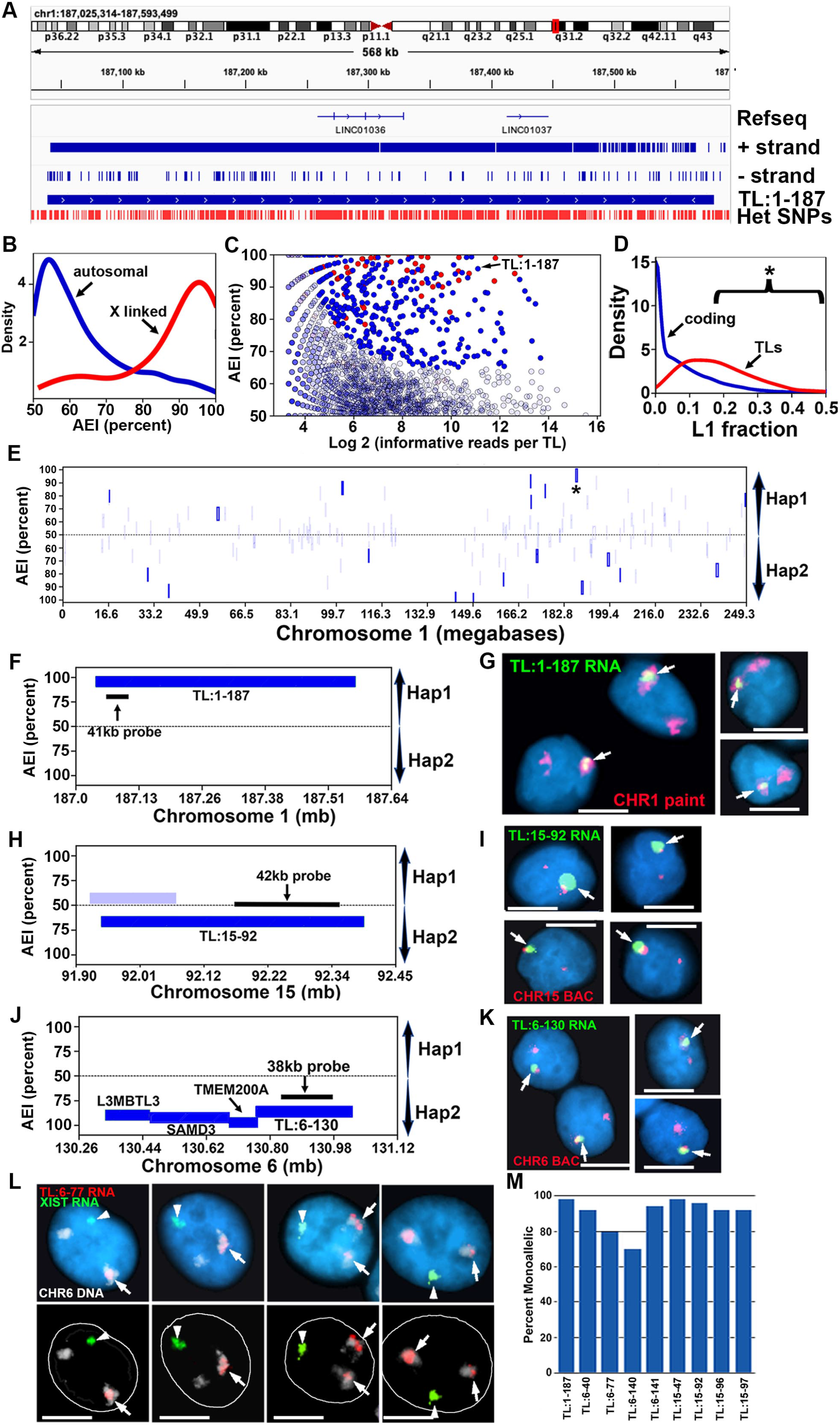
Genome wide allelic expression imbalance and spatial retention patterns of TL in GM12878. A) Genome browser view of a representative example of an intergenic non-coding contiguously transcribed locus on the plus strand of chromosome 1. The location of 562 heterozygous SNPs within TL:1-187 are shown in red along the bottom of the panel. B) Genome wide distribution of AEI of TLs on all autosomes (blue) and the X chromosome (red). We define AEI as an allelic-bias that is outlying parametric (FDR-Benjamini Hochberg corrected binomial test q-value<=0.01) and non-parametric (allelic-bias >=2.5 s.d. above autosome-wide mean, controlled for expression level) estimation of the null-distribution of bi-allelic expression. C) Scatter plot of AEI of autosomal TL (blue dots) and X-linked TL (red dots) as a function of the number of informative reads. Opaque dots are outliers on the genome-wide distribution of AEI. D) Distribution of the fraction of L1 derived sequence in TL and coding genes. E) Chromosome 1 view of AEI of TL (opaque: statistically significant). The position of TL:1-187 is shown. F, H, J) Zoom-in views of AEI of representative TL (Dark blue = statistically significant; light blue not significant). The location and size of Fosmid probes (see Table S2) used for RNA FISH are shown. G, I, K) RNA-DNA FISH images of TL:1-187, TL:15-92, and TL:6-130 expression in single cells. L) RNA-DNA FISH image of TL:6-77 RNA (red), CHR6 DNA (chromosome paint, white), and XIST RNA (green) visualized within individual cells, top and bottom panels represent the same four cells with the nuclear outline drawn in white. M) Percent of cells exhibiting strong AEI (Percent Mono-allelic) for nine representative TLs, as measured by DNA/RNA FISH visualization.

**Table1.** Summary of autosomal AEI and VERT loci.

| **Sample** | **TL** | **AEI TLs** | **AEI Coding** | **Random AEI TL** | **Random AEI Coding** | **VERT loci** | **AEI TL within VERT** | **AEI coding within VERT** |
| --- | --- | --- | --- | --- | --- | --- | --- | --- |
| EB3_2<br>(6 clones) | 2,692 | 644<br>(24%) | 941<br>(12.5%) | 68 | 62 | 99<br>(68mb) | 82 | 80 |
| GM12878<br>(parent) | 1,492 | 300<br>(20%) | 380<br>(5%) | N/A | N/A | N/A | N/A | N/A |
| GM12878<br>(2 clones) | 1,815 | 378<br>(21%) | 613<br>(8.5%) | 6 | 4 | 139<br>(124mb) | 44 | 57 |

To determine if TL RNAs with AEI remain associated with their parent chromosomes, we used RNA-DNA FISH to assay for chromosome territory localized retention of TL:1-187 plus 8 additional TL in GM12878 cells. For this analysis we used Fosmid probes to detect TL RNA (Fig. 1F, 1H, and 1J; and see Table S2) and a chromosome-specific paint, BAC (Bacterial Artificial Chromosome) or centromeric probe to detect DNA for the chromosome of interest. We detected RNA hybridization signals that remain associated with their respective chromosomes for all 9 TL RNAs (for examples see Figs. 1G, 1I, 1K, and 1L), which is consistent with the spatial-expression pattern observed for the known ASARs ^4-6^. For an independent assessment of mono- and bi-allelic expression, we quantified the frequency of single and double sites of RNA hybridization. For this analysis we included an XIST RNA FISH probe as positive control (GM12878 have a single large site of RNA hybridization signal for XIST in >90% of cells), and scoring cells with a single XIST RNA FISH signal and TL RNA hybridization signals indicated that >70% of cells had single sites of hybridization for each TL (Fig. 1M). Two sites of RNA hybridization were detected for all 9 TL, ranging from 2% to 30% of cells (see Fig. 1L for examples). We note that the size of the RNA hybridization signals detected by all TL probes was variable, ranging in size from large clouds that occupy the entire chromosome territory to relatively small sites of hybridization.

The observations described above indicate that ∼300 autosomal TL are expressed in an allele restricted manner in GM12878. We next sought to determine if the AEI of TL was present in LCLs from a second unrelated individual, and if the AEI could come from either allele in multiple single-cell derived clones, which would distinguish between random epigenetically programed AEI from imprinted AEI and AEI caused by genetic polymorphisms (*i. e*. eQTL). For this analysis, we used EB3_2 LCLs, which were isolated from a female with a haplotype-phased genome ^46^. By leveraging the B-cell origin of LCLs, we isolated six single-cell derived clones, as defined by the presence of unique immunoglobulin gene rearrangements [see Table S4; ^6,21,47^]. The EB3_2 clones (EB2, EB3, EB4, EB10, EB13, EB15) are expected to be isogenic except at the regions of immune gene related somatic rearrangements, and were expanded for >25 population doublings prior to the generation of nuclear enriched RNAseq and Repli-seq libraries (see Fig. 2A). Because EB3_2 cells are female, we first queried AEI on the X chromosome. The six EB clones display AEI of 66 TL and 127 protein coding genes located on the X chromosome (Table S1). The orientation of the expressed alleles indicated that all six EB clones have the same active and inactive X chromosomes, haplotype 1 (maternal) and haplotype 2 (paternal), respectively. Figure 2B shows examples of the AEI of X chromosome TL and protein coding genes, and indicates strong expression from haplotype 1 in all 6 clones.

**Figure 2.**
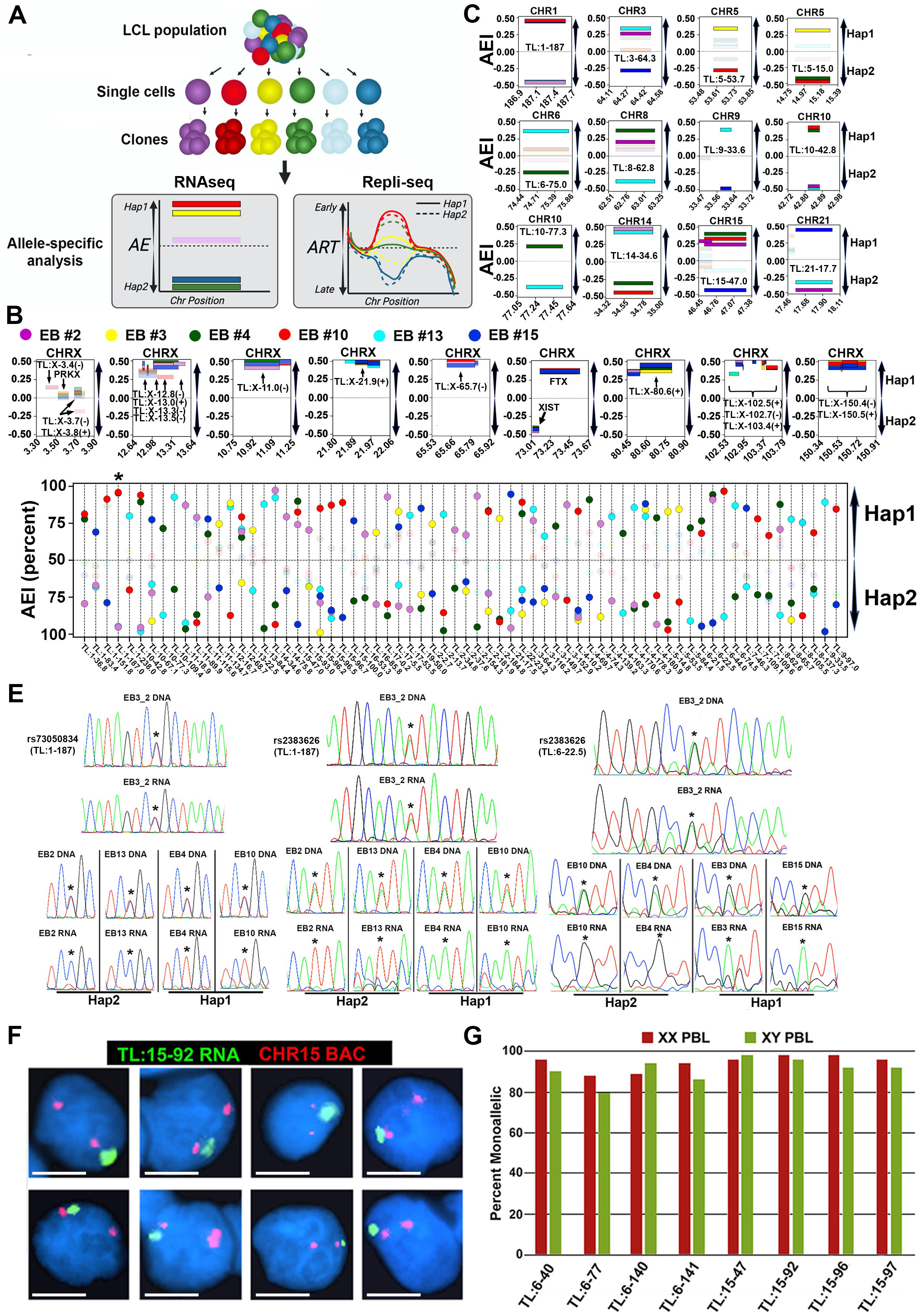
Haplotype resolved analysis of allelic expression imbalance of TL RNAs. A) Sub-cloning and allele-specific genomic analysis scheme for 6 clones from the EB3_2 lymphoblastoid cell line. The left panel shows phased RNAseq data at one hypothetical TL showing AEI in different clones. The right panel shows the phased Repli-seq data at one hypothetical replication domain showing synchronous or asynchronous replication. B-C) Zoom-in views of examples of TL RNAs that display AEI (Hap1: haplotype 1, Hap2: haplotype 2) on the Y axis and the genomic positions (in megabases) for X chromosome (CHRX; panel B) or autosome (panel C) positions are shown on the X axis. TL and protein coding genes are labeled and marked by arrows. C) Each panel represents the AEI and location of a prominent TL from 9 different autosomes (CHR1 etc). D) All TL RNAs that display AEI from different alleles within the 6 EB3_2 clones (X-axis: chromosome start-position; Y-axis: AEI as percent). E) Sanger sequencing traces of PCR products generated from genomic DNA and cDNA illustrating heterozygosity in genomic DNA and mono-allelic expression of either allele from TL RNAs (EB3_2: parent cell population, EB-2, 3, 4, 10, 13, and 15: clones). F) RNA-DNA FISH in PBLS using a Fosmid probe to TL:15-92 to detect RNA and a chromosome 15 BAC to detect DNA. G) Quantification of the number of RNA FISH signals indicated that 8 different TL were detected from single sites in >80% of cells in PBL populations isolated from two (one female XX and one male XY) unrelated individuals. We also note that 2%-20% of cells contained two sites of hybridization.

By analyzing expression from autosomes across all six clones, 2,692 TL were defined in the EB3_2 clone set, with 644 TL, or ∼24%, showing statistically significant AEI (Table 1). Strikingly, 68 TL were identified that display AEI from opposite homologs in two or more clones (Fig. 2C and 2D), which is consistent with random epigenetically programed AEI. In addition, some clones display equivalent levels of expression between alleles (*i. e*. bi-allelic expression) at the same TL that display AEI of either allele in other clones, while in other clones there is undetectable expression. The other 576 autosomal TL with AEI showed expression from the same allele, both alleles or neither allele in one or more clones (Table S1). This AEI pattern is not consistent with genomic imprinting, nor with the presence of eQTL, but may be due to random epigenetically programed AEI where we have not analyzed enough clones to detect expression from opposite alleles. Regardless, validation of the clone-specific AEI from autosomal TL was then performed by sanger sequencing of DNA and cDNA from the parent multiclonal population and from individual clones (see Fig. 2E for examples). These data indicate that expression of 68 autosomal TL can originate from either haplotype 1, haplotype 2, both or neither in different clones isolated from the same individual. These observations indicate that each allele had acquired the expressed or silent state independently from the other allele, and because all possible combinations were detected in different clones from the same individual these differential expression states must be under epigenetic control.

Next, to determine if the TL RNAs that show AEI in LCLs are associated with their parent chromosome in human primary cells, we carried out RNA-DNA FISH using probes to 8 different TL on primary blood lymphocytes (PBLs) isolated from two unrelated individuals. We detected chromosome-associated RNA hybridization signals for all 8 TL. An example of this analysis is shown for TL:15-92 in Figure 2E. Quantification of the number of RNA FISH signals in >100 cells indicated that all 8 TL were expressed from single chromosomes in >80% of cells (Fig. 2G). We note that the size of the RNA hybridization signals detected by all of the TL probes was variable, ranging in size from large clouds to relatively small sites of hybridization. We also detected two sites of hybridization for all TL probes in 2% to 20% of cells (Fig. 2G), which is consistent with the LCL RNAseq data (see Fig. 2B and 2C), and indicates that expression and chromosome territory localization of this set of TL RNAs can be bi-allelic in at least some cells in the PBL populations. Therefore, we conclude that mono- and bi-allelic expression of chromosome associated RNAs can be detected for all 8 TL in primary cells.

### Variable Epigenetic Replication Timing occurs at hundreds of autosomal loci

ASRT occurs at regions of autosomes that display epigenetically programmed AEI ^13,28-32^. To assess the relationship between AEI and ASRT in LCLs, we next performed allele-specific Repli-seq on the six EB3_2 clones (see Fig. 2A), and on two single cell derived clones from GM12878. First, allele-specific replication timing profiles were generated for each haplotype from each clone from both sets of clones, totaling 12 and 4 allele-specific RT profiles per chromosome for the EB3_2 and GM12878 clone sets, respectively. Second, differences in RT among each clone set was assessed by measuring the standard deviation (SD) of three subdivisions of the RT profiles. For each clone set we generated a “combined RT profile” that included haplotype 1 plus haplotype 2 (e. g. all 12 alleles in the six EB3_2 clones); and two “allele-specific RT profiles” that included only haplotype 1, or only haplotype 2. Third, to identify classical ASRT regions within individual clones, we analyzed the difference between haplotype 1 and haplotype 2 RT profiles within each clone separately by comparing the absolute difference in allele-specific read counts in the Repli-seq data, which can detect ASRT resulting from either genetic or epigenetic mechanisms ^3^. Figure 3A illustrates hypothetical RT profiles for a region of the genome that shows synchronous or asynchronous RT, as well as epigenetic variability that could occur in the RT profiles in different clones from the same individual.

**Figure 3.**
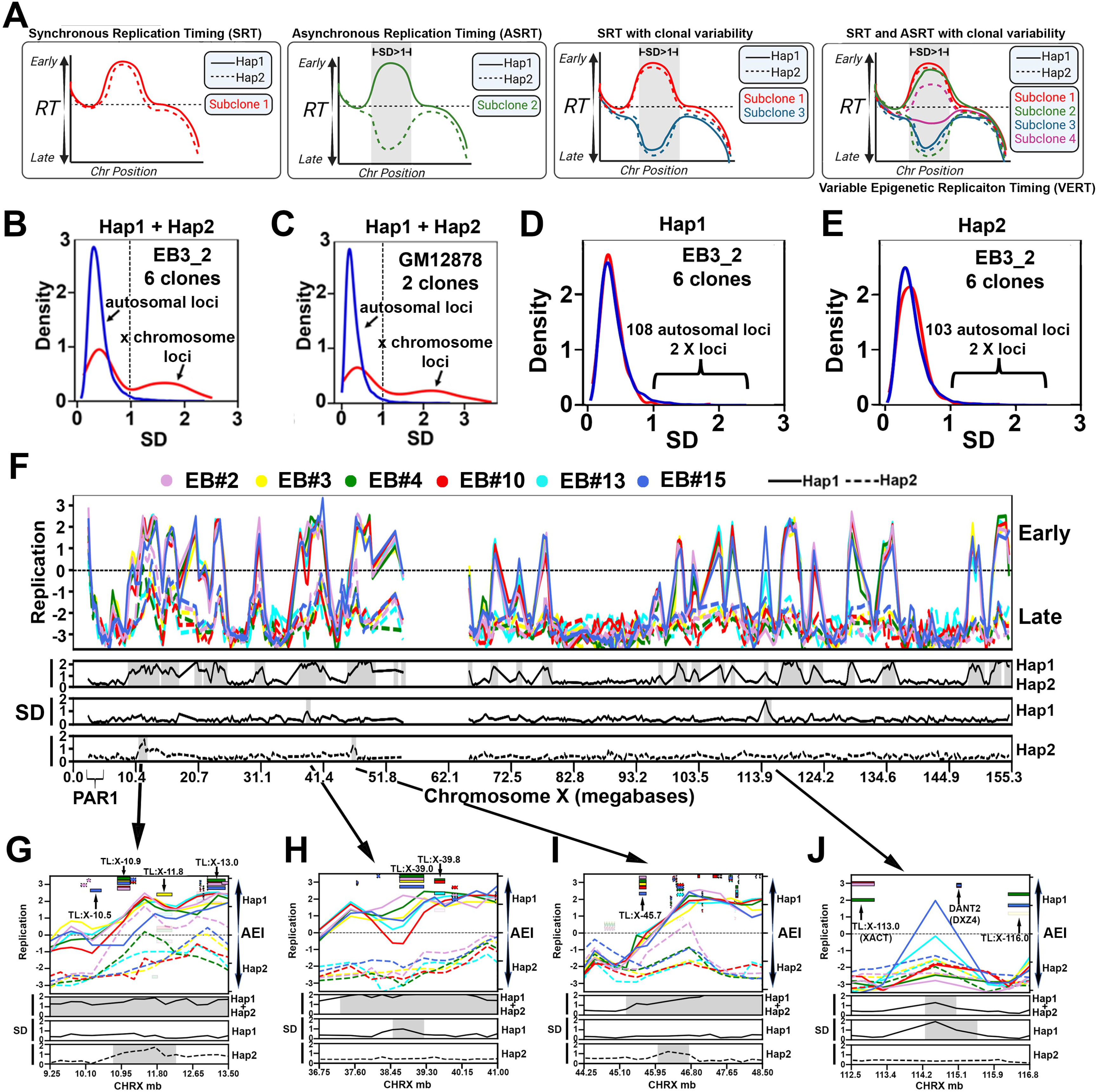
Haplotype resolved analysis of asynchronous replication timing. A) Illustration of a hypothetical genomic region that shows Synchronous Replication Timing (SRT), Asynchronous Replication Timing (ASRT), and possible clonal variability in both SRT and ASRT. Areas with standard deviation (SD) >1 are highlighted in gray. B and C) Genome-wide distribution of the SD of replication timing of individual alleles among clones derived from EB3_2 (12 alleles) and GM12878 (4 alleles). Standard deviation of RT between alleles was calculated for each 250kb genomic window, generating a distribution with median of 0.37 and standard deviation of 0.23 RT units (Early/Late Log2Ratio) in EB3_2, and median 0.25 with standard deviation of 0.26 in GM12878, indicating higher similarity between the two GM12878 subclones. Outliers were identified as: SD ≥ 2.5x SD + mean: >0.92 for the EB3_2 clones and >0.90 for the GM12878 clones. D and E) Genome-wide distribution of the standard deviation (SD) of replication timing of each individual allele among clones derived from EB3_2 (D: 6 Hap1 alleles; E: 6 Hap2 alleles). Outliers were identified as: SD ≥ 2.5x SD + mean: >0.96 for Hap 1 and >0.98 for Hap 2 in the EB3_2 clones and >0.92 for Hap 1 and >0.94 for Hap 2 in the GM12878 clones. F) Chromosome X Early/Late RT profile with SD of haplotype-resolved replication timing from the 6 EB3_2 clones highlighting (gray shading) outlier regions from the “combined RT profile” (Hap1 plus Hap2) and from the “allele-restricted RT profile” separately (Hap1 or Hap2). Each clone was color coded as shown, with haplotype 1 shown as a solid line, and haplotype 2 shown as a dotted line for both sets of clones. The left axis shows the RT (Early/Late) profiles, with positive numbers representing early replication and negative numbers representing late replication. G-J) Zoom in views of the 4 locations on the X chromosome that contain outliers in SD of the “allele-restricted RT profiles”. Each clone was color coded as shown, with haplotype 1 shown as a solid line, and haplotype 2 shown as a dotted line for both sets of clones. The left Y axis shows the Early/Late RT profiles. The SD of the replication timing across each locus is shown below each panel. Areas highlighted in gray represent outliers in the SD. The right Y axis shows the AEI, with smooth rectangles representing TL and the stippled rectangles representing protein coding genes. The opaque rectangles show AEI (FDR-BH alpha <=0.01, while the transparent rectangles show bi-allelic expression (FDR-BH alpha <=0.01) of TL and protein coding regions.

The genome wide distribution of the SD for the “combined RT profile” on autosomes for each 250kb genomic region revealed a single large peak (SD <1) of synchronous replication with a shallow but long right tail of outliers (SD ≥ 2.5 x SD + mean: >0.92 for the EB3_2 clones and >0.90 for the GM12878 clones; Fig. 3B and 3C). In contrast, the distribution of the SD of the “combined RT profile” along the X chromosome revealed two peaks. One peak is superimposed with the synchronous autosomal peak (SD <1), and these X chromosome regions represent the late replicating regions on both the active and inactive X chromosomes (Fig. 3B, 3C and 3F). The X chromosome regions within the second broader peak (SD >1) map to the regions of the X chromosome that has early replication (Fig. 3F), and the early replicating DNA is from haplotype 1, which represents the active X chromosome in all six EB3_2 clones (see Fig. 2B and Table S1). The “combined RT profile” analysis of the two GM12878 clones revealed a similar profile for the autosomes and X chromosomes (Fig. 3C), with haplotype 2 representing the early replicating active X chromosome in both clones (see Table S1).

For the “allele-restricted RT profiles”, we compared the RT profiles of each haplotype in all 6 EB3_2 and both GM12878 clones independently from the RT profiles of the other haplotype in each clone set (Fig. 3D and 3E). This analysis allowed us to detect outliers in the RT profiles on each allele in the two clone sets separately (SD ≥ 2.5 x SD + mean: >0.96 for Hap 1 and >0.98 for Hap 2 in the EB3_2 clones; and >0.92 for Hap 1 and >0.94 for Hap 2 in the GM12878 clones). In contrast to the “combine RT profile” analysis described above (Fig. 3B and 3C), the “allele-restricted RT profile” analysis identified only four relatively small outlier regions on the X chromosome, two on the active and two on the inactive X chromosomes (Fig. 3F-3J). All four of these regions contain TL with strong AEI (Fig. 3G-3J), and one of these regions contains two loci, XACT and DXZ4, that are known to be important for X inactivation in humans [see Fig. 3J; and ^48-51^]. Because this Repli-seq analysis was performed on clones isolated from the same individual and the differences in replication timing between the active and inactive X chromosomes are known to be under epigenetic control, we conclude that this analysis allows us to detect epigenetically controlled allele-specific replication timing.

The “combined RT profile” analysis indicated that the vast majority of autosomal DNA replicated synchronously (SD <1), with a relatively small number of loci (99 loci in the EB3_2 and 139 loci in the GM12878 clone sets) with significant differences in allele-specific RT (Table S3). In addition, analyzing the “allele-restricted RT profiles”, we identified 108 haplotype 1 and 103 haplotype 2 outlier loci in the EB3_2 clones (Fig. 3D and 3E). In total, we identified outliers in the autosomal RT profiles for 248 loci in the EB3_2 plus GM12879 clones, with 37 loci shared between the two clone sets (211 loci representing ∼190 mb or ∼6% of the human genome; Table 1). Figure 4A shows the RT profiles on chromosome 5 in the six EB3_2 clones, and indicates that there are 9 loci in the “combined RT profile” with significant differences in RT, and the “allele-restricted RT profiles” indicated that these differences are associated with either haplotype 1, haplotype 2 or both. Figure 4B-4I shows examples of 8 loci identified as outliers in both EB3_2 and GM12878 RT profiles. The differences in the “allele-restricted RT” profiles detected at outlier loci can affect either haplotype 1, haplotype 2 or both. Thus, some clones display large differences in RT between alleles at the same locus that displays synchronous replication in other clones, and the RT profiles in the synchronous clones can either have the earlier or later replication timing profile (Fig. 4B-4I; also see Fig. 5B-5J and Fig. 3A). Taken together, these observations indicate that each allele had acquired the earlier or later replication independently from the other allele. Because the differences in RT at these loci are quite variable and affect either allele independently, we refer to these loci as having <u>V</u>ariable <u>E</u>pigenetic <u>R</u>eplication <u>T</u>iming (VERT). Within the 99 VERT loci in the EB3_2 clone set we identified 82 TL that display AEI (Table 1 and Table S1 and S3; also see Fig. 4B-I and Fig. 5 below). In addition, at any given TL with AEI the expression could come from either the earlier or later replicating allele, indicating that each allele had acquired the expressed or silent state independent of the replication timing within these VERT loci.

**Figure 4.**
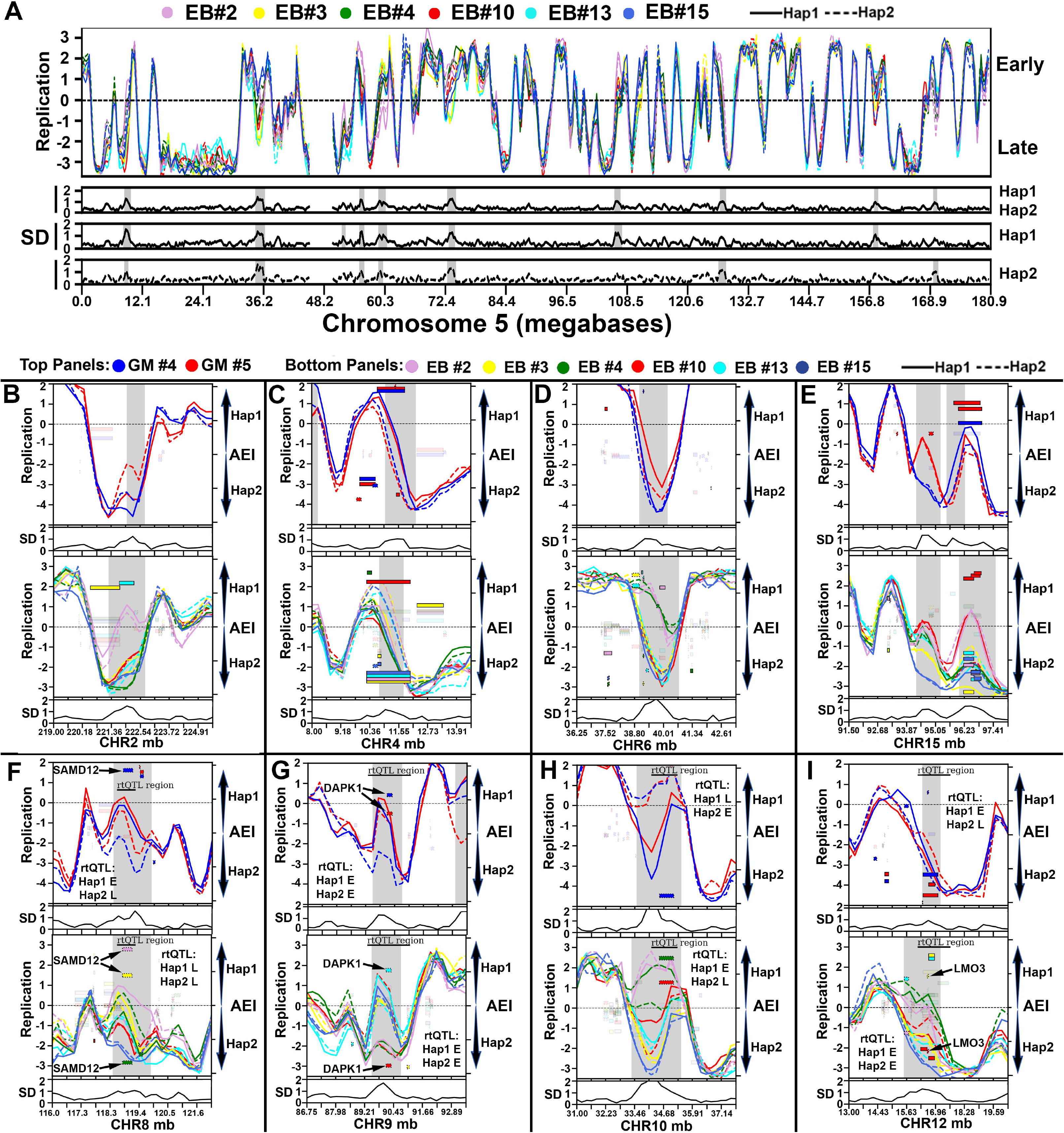
Haplotype phased analysis of ASRT on autosomes. A) Chromosome 5 RT profile from the 6 EB3_2 clones, highlighting regions with SD >1 from both the “combined allele RT profile” (Hap1 and Hap2) and from the “allele-restricted RT profile” separately (Hap1 or Hap2). B-I) Representative examples of regions present in both GM12878 (top panels) and EB3_2 (bottom panels) clones with variable epigenetic replication timing between alleles. Each clone was color coded as shown, with haplotype 1 shown as a solid line, and haplotype 2 shown as a dotted line for both sets of clones. The left Y axis shows the replication timing (Early/Late) profiles. The SD of the replication timing across each locus is shown below each panel. The right Y axis shows the AEI, with smooth rectangles representing TL and the stippled rectangles representing protein coding genes. The opaque rectangles show AEI (FDR-BH alpha <=0.01), while the transparent rectangles show bi-allelic expression of TL and protein coding regions. Areas highlighted in gray represent outliers in the SD. F-I) The location of rtQTL at VERT regions, with the Early (E) and Late (L) alleles ^26^ for both GM12878 and EB3_2 shown (also see Table S5).

**Figure 5.**
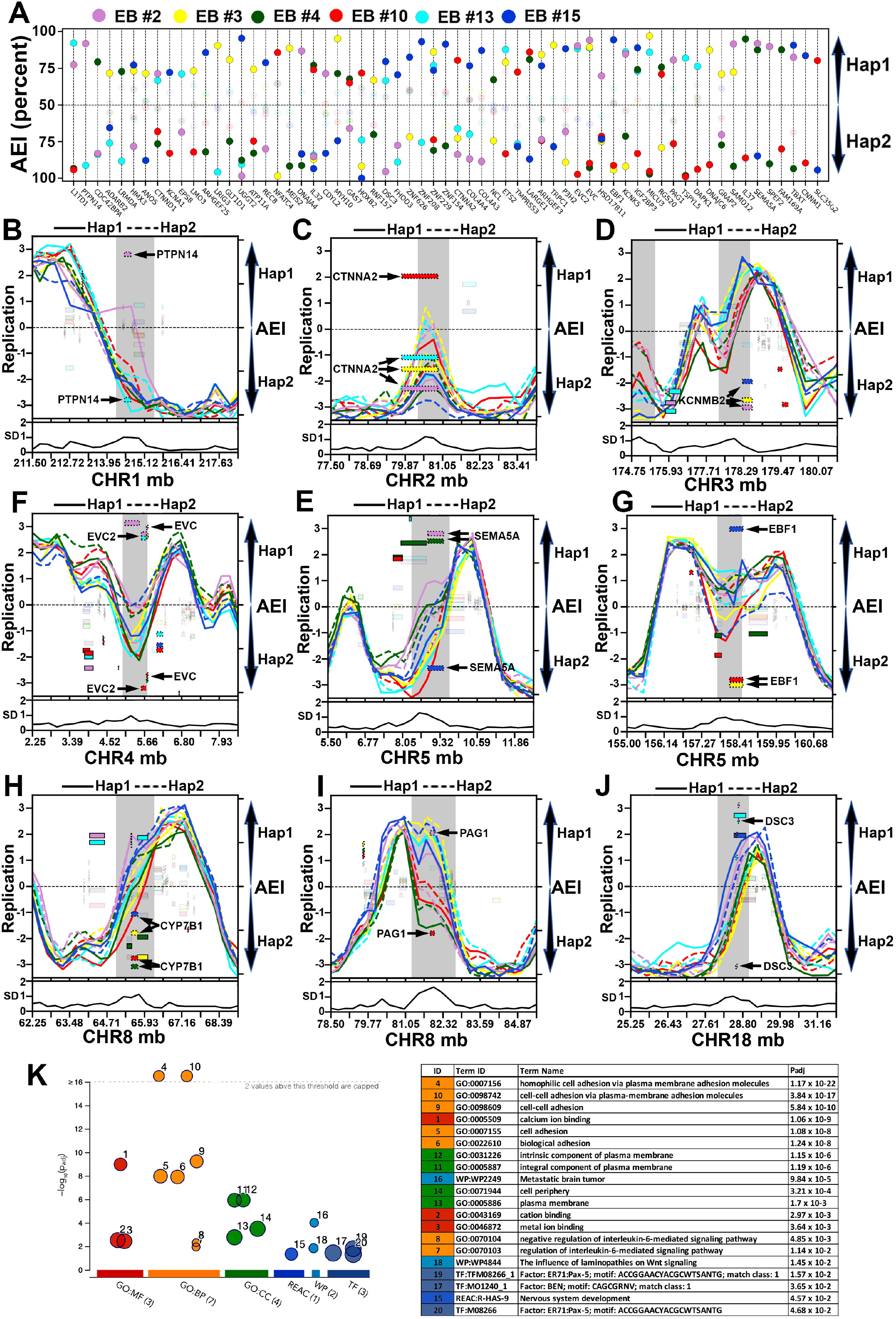
Haplotype resolved expression and replication of protein-coding genes. A) 63 protein coding genes display random epigenetic AEI in EB3_2 clones. (X-axis: protein coding gene; Y-axis: AEI). B-J) Examples of genomic regions that contain protein coding genes that display epigenetic AEI and VERT in the six EB3_2 clones. The left axis shows the replication timing (Early/Late) profiles, and the standard deviation (SD) across each locus is shown below each panel. The right axis shows the AEI, with the stippled rectangles representing protein coding genes and smooth rectangles representing TL. The opaque rectangles show AEI (FDR-BH alpha <=0.01), while the transparent rectangles show bi-allelic expression of protein coding and non-coding transcripts. K) Functional enrichment analysis of coding genes that display random epigenetic AEI in EB3_2 clones.

### Asynchronous replication can occur at rtQTL, and lacks coordination between loci

By performing allele-specific Repli-seq on single-cell derived clones, we were able to address the consequences of DNA sequence polymorphisms on ASRT. A previous study identified 20 polymorphic loci, known as rtQTL, by examining RT profiles in human LCLs ^26^. We found that 12 of the 20 previously identified rtQTL were within VERT loci detected in either the EB3_2, GM12878, or both Repli-seq data sets. Surprisingly, we found that these 12 loci display strong epigenetic differences in replication timing regardless of which rtQTL allele they contain (Table S5). Examples of 4 different VERT loci that were detected in both GM12878 and EB3_2 clones, with the genotype and location of the associated rtQTL, are shown in Figure 4F-4I (also see Fig. S3A). We found that differences in RT within these VERT regions occur at loci that are either homozygous or heterozygous for the rtQTL alleles, demonstrating that the epigenetic effects described here are dominant over any DNA sequence polymorphisms that are present at these loci.

We next sought to address the question of coordination of asynchronous replication timing. We and others previously reported that ASRT of random mono-allelic genes is coordinated with other random mono-allelic genes on the same chromosome, suggesting that there is a chromosome-wide system that coordinates replication asynchrony of random AEI genes ^4,9,28,32,52^. More recently, the asynchrony associated with random AEI genes in mouse pre-B cells was proposed to be coordinated on all autosomes, resulting in only two “mirror-image” patterns of asynchronous replication ^53,54^. For the analysis of coordination at ASRT regions utilizing Repli-seq, we first defined ASRT regions as outliers on the distribution (SD >1) of the difference between haplotype 1 and haplotype 2 RT profiles within each EB3_2 clone. We then classified each allele at each asynchronous region as either early or late within each clone, and compared the Early/Late status across multiple regions on individual chromosomes and between clones. We first analyzed the ASRT on the X chromosome (Figure S1A). We detected dramatic coordination in the polarity of the ASRT regions along the length of the X chromosome in all six clones (see Fig. S1), which is consistent with previously reported ASRT profiles for the active and inactive X chromosomes in LCLs ^55^. While we detected numerous loci on every autosome that display ASRT, we found no evidence for coordination, either on the same autosome or between autosomes (see Fig. S1B-D). Furthermore, by analyzing the Repli-seq data previously published by Blumenfeld et al for mouse pre-B cell clones ^53^ using our “combined RT profile” and ASRT analyses described here, we detected loci in the mouse genome that display VERT, but did not detect coordination in the Early/Late replication timing pattern at loci that show ASRT, and therefore we do not detect a “mirror image” coordination pattern on mouse chromosomes in the Repli-seq data for mouse pre-B cell clones (Fig. S2). Our data indicate that the clonal Early/Late pattern at multiple asynchronous loci on human or mouse autosomes is not coordinated on the same chromosome nor between different chromosomes.

### Variable epigenetic AEI of protein coding genes occurs at VERT loci

The observations described above indicate that hundreds of TL exhibit AEI, and that this AEI often occurs at genomic loci detected as having VERT. We next determined if protein coding genes also display epigenetically programmed AEI within VERT loci. Utilizing allele-specific analysis of the RNAseq data at protein coding genes, we identified 941 protein coding genes that display AEI, including 11 known imprinted genes (geneimprint.com; Table S1). We also identified 61 protein coding genes that display AEI from opposite homologs in two or more clones (Fig. 5A and Table 1), which is consistent with random epigenetically programed AEI. Similar to the TL with epigenetically programed AEI, some clones display equivalent levels of expression between alleles (*i. e*. bi-allelic expression) while other clones express only one allele, and in yet other clones have undetectable expression. These observations are consistent with previous work that found a similar variability in mono-allelic expression of protein coding genes within LCL clones ^21^. We found that 98 of the 941 protein coding genes with AEI are within VERT regions (Table 1; also see Tables S1 and S3). Figure 5B-J shows examples of the RT profiles and AEI of protein coding genes at genomic regions that also display VERT (also see Fig. 4F, 4G, and 4I). These observations indicate that the expressed or silent state at these protein coding genes was acquired independently, and that the expressed or silent state is not associated with either earlier or later replication. Utilizing an ensemble of gene enrichment analysis tools ^56^, the 61 protein coding genes with random epigenetically programed AEI were found to be enriched for several biological processes, with homophilic cell adhesion being the most significant (P=1.17×10E^-22^; Fig. 4K).

### Deletion of TL within VERT loci result in delayed replication timing in *cis*

The previous ASAR genetic disruption studies were carried out in cells that displayed strong AEI at the ASAR loci, and disruptions of the expressed alleles resulted in delayed replication while disruptions on the silent alleles did not ^4-6,10^. However, the variability in the AEI described in this report indicates that TL can be expressed from both alleles in at least some clones, and that the expressed alleles are independent of earlier or later replication at the locus. To determine if TL that reside in VERT loci control replication timing of entire chromosomes, we used CRISPR/Cas9 to delete 5 different TL in HTD114 cells. We chose HTD114 cells for this analysis because they contain AEI and ASRT of both imprinted and random mono-allelic genes, they have a stable karyotype that does not change significantly following transfection, drug selection and sub-cloning, and knockouts of the three known ASAR genes result in delayed replication timing, delayed mitotic condensation and chromosome structure instability in *cis* ^4-10^. However, given the tissue-restricted expression pattern of TL, we first determined which of the lymphocyte TL with AEI and VERT are expressed in HTD114 cells. We first queried RNAseq data from HTD114 cells [see ^5^], and subsequently used RNA-DNA FISH and Sanger sequencing of PCR products from reverse transcribed RNA for 16 TL to determine if the RNAs are localized to their parent chromosomes and whether or not they show mono-versus bi-allelic expression. All 16 TL RNAs were found to be associated with their parent chromosomes, and displayed single or double RNA FISH signals in the nuclei of HTD114 cells (for examples see Fig. 6B-6C, Fig. S3B-S3C, and S4B-S4C). Because the epigenetic variability that results in AEI at TL can result in mono- or bi-allelic expression in individual clones, we chose TL that were either mono- or bi-allelic in the HTD114 cells for genetic deletion assays. We chose 5 TL, two expressed from both alleles, TL:1-187 (Figure 6) and TL:8-2.7 (Figure S3), and three expressed from single alleles TL:9-23, TL:9-24, and TL:9-30 (Figure S4), for CRISPR/Cas9 mediated deletions.

**Figure 6.**
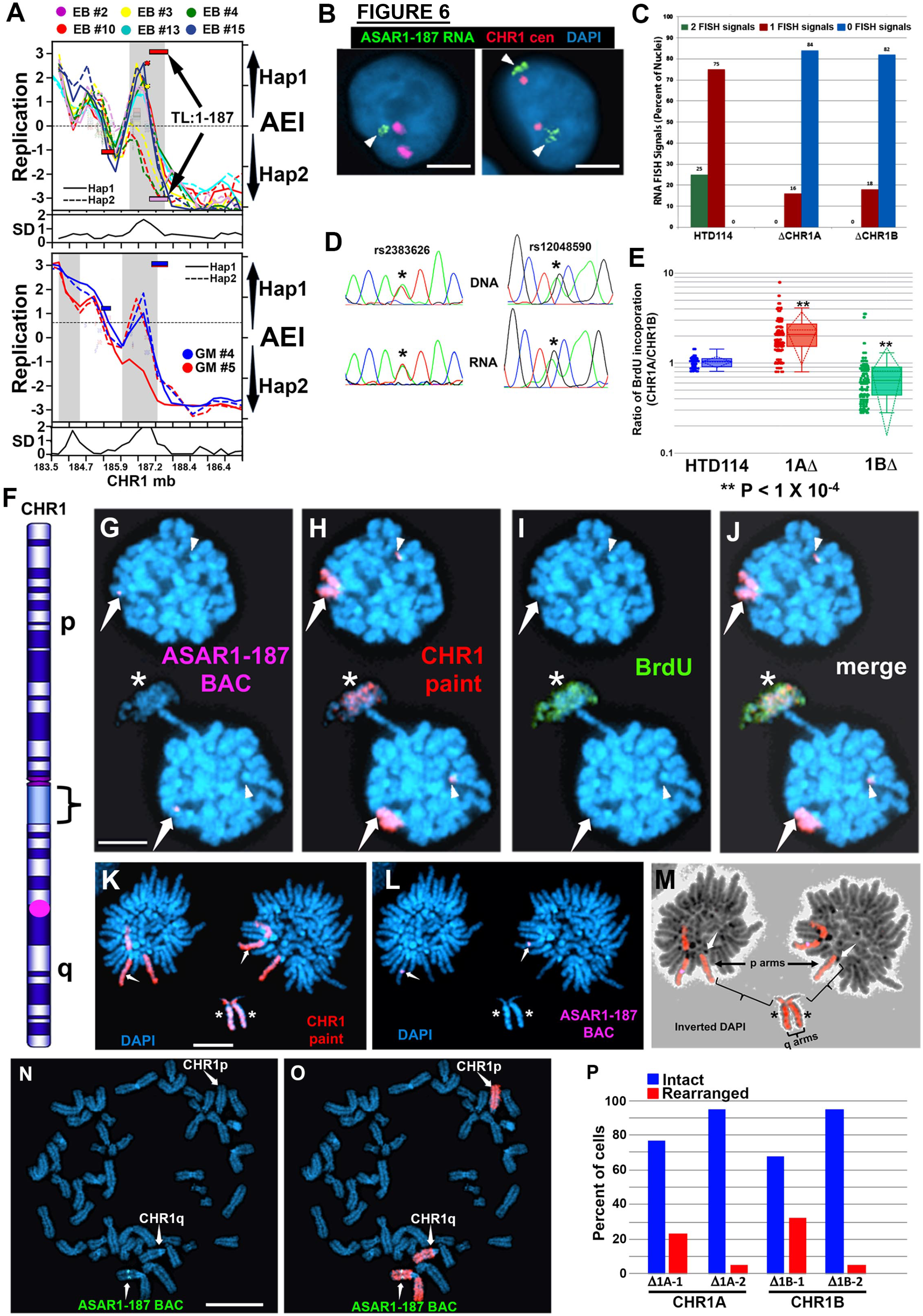
Delayed replication of chromosome 1 following disruption of TL:1-187. A) AEI and ASRT at the TL:1-187 locus in the EB3_2 (top panel) and GM12878 (bottom panel) clones. The left axis shows the replication timing (Early/Late) profiles, and the standard deviation (SD) across the locus is shown below each panel. The right axis shows the AEI of haplotype 1 (Hap1) and haplotype 2 (Hap2). The smooth rectangles represent TL, with TL:1-187 marked with arrows. The stippled rectangles represent protein coding genes. The opaque rectangles show AEI (FDR-BH alpha <=0.01), and the transparent rectangles show bi-allelic expression of protein coding and non-coding transcripts. B) Examples of cells analyzed using RNA-DNA FISH using at TL:1-187 probe (green; see Table S2) to detect RNA and a chromosome 1 centromeric (CHR1 cen) probe to detect DNA. Arrowheads mark the sites of TL:1-187 RNA FISH hybridization. C) Quantitation of the RNA FISH hybridization signals for TL:1-187 before and after allele-specific deletion of the locus in the HTD114 cell line. D) Sanger sequencing traces of PCR products generated from HTD114 genomic DNA and cDNA illustrating heterozygosity, at two SNPs (rs2383626 and rs12048590) within TL:1-187. Note that both SNPs are heterozygous in genomic DNA and heterozygous in the cDNA samples. E) BrdU incorporation in HTD114 cells containing heterozygous deletions of the TL:1-2.7 locus. Cells were processed for BrdU incorporation and processed for DNA FISH using a chromosome 1 centromeric probe plus a BAC from within the deleted region (see panel F purple dot; and Table S2). The TL:1-187 BAC was used to identify the chromosome 1 with the deletion (see G-N). Quantification of BrdU incorporation in multiple cells with heterozygous deletions indicated that deletion of either allele resulted in delayed replication of chromosome 1 (1AΔ or 1BΔ). F) Illustration of chromosome 1 with the location of the short (p) and long (q) arms, G-banding pattern, the large heterochromatic block (bracket), and the location of TL:1-187 (purple dot). G-J) Delayed replication timing and delayed mitotic chromosome condensation on chromosome 1 in HTD114 after deletion of the TL:1-187 locus. Cells were processed for BrdU incorporation (green) and for DNA FISH using a chromosome 1 paint probe (red) plus a BAC from within the TL:1-187 deleted region (purple; see panel F). The TL:1-187 BAC was used to identify the chromosome 1 with the deletion. An anaphase cell containing a lagging chromosome 1 is shown. The arrows mark chromosome 1s that retain the TL:1-187 BAC hybridization signal, and the chromosome 1 paint hybridized to the lagging chromosome, which also lacked a detectable TL:1-187 BAC signal (asterisk). In addition, the only chromosome in this cell with BrdU incorporation was the lagging chromosome 1 (asterisk). We also detected a fragment of chromosome 1 translocated to an unknown chromosome (arrowheads) that was present in both sets of anaphase chromosomes. K-M) An anaphase cell with a lagging chromosome 1 is shown. This cell was processed for DNA FISH using a chromosome 1 paint probe (red) plus a BAC from within the TL:1-187 deleted region (purple; see panel F). Note that the lagging chromosome lacks the TL:1-187 hybridization signal on the q arm (asterisks). The arrows mark the chromosome 1s that retain the TL:1-187 BAC hybridization signal, and the chromosome 1 paint hybridized to the lagging chromosome, which also lacked a detectable TL:1-187 BAC signal. Also note the extremely under-condensed region associated with the large heterochromatic block on the long arm of chromosome 1, which is highlighted with brackets in the inverted DAPI image shown in panel M. The short (p) and long (q) arms of the lagging chromosome 1 are shown, and the arrows in panel M mark the centromeres of the lagging chromosome 1s. N, O) Chromosome 1 rearrangements are common following TL:1-187 deletion. A mitotic spread is shown that was processed for DNA FISH using the TL:1-187 BAC (green) and a chromosome 1 paint (red). Note the broken chromosome 1 lacks the TL:1-187 BAC signal. P) Quantitation of the chromosome 1 rearrangements in heterozygous deletions of TL:1-187. All rearrangements of chromosome 1 that we detected affected the deleted chromosome, which was either CHR1A or CHR1B.

TL:1-187 is expressed from chromosome 1 at ∼187-187.5 mb (see Fig. 1A, 1E-G; and 2C), and Figure 6A shows the AEI and RT profiles from the EB3_2 and GM12878 clone sets. Figure 6B and 6C shows results from the RNA-DNA FISH analysis in HTD114 cells, and indicates that single sites of RNA hybridization were detected in ∼75% of cells and that two RNA hybridization signals were detected in ∼25% of cells, suggesting that HTD114 cells express both alleles. Figure 6C shows sequencing traces from two different SNPs (see Table S4) that are heterozygous in genomic DNA, and heterozygous in reversed transcribed RNA isolated from HTD114 cells, confirming that the TL1-187 transcripts are generated from both alleles in HTD114 cells.

Next, we designed sgRNAs to unique sequences flanking the genomic region that expresses TL:1-187 (Table S4). We expressed these sgRNAs in combination with Cas9 and screened clones for deletions using PCR primers that flank the sgRNA binding sites (see Table S4). Because TL:1-187 expression is bi-allelic in HTD114 cells, we isolated clones that had heterozygous deletions affecting either homolog. We determined which allele was deleted based on retention of the different base pairs of heterozygous SNPs located within the deleted regions, and arbitrarily assigned the homologs as CHR1A or CHR1B (see Fig. 6D and Table S4). In addition, to determine how deletion of each allele affected the number of RNA hybridization signals, we carried out RNA-DNA FISH using the TL:1-187 probe in heterozygous deletion clones. Figure 6C shows the quantification of the RNA FISH signals, and indicates that deletions of TL:1-187 from either CHR1A or CHR1B resulted in cells with single sites of RNA hybridization signals, and no cells with two sites of hybridization, indicating that both alleles of TL:1-187 are expressed in HTD114 cells.

We next assayed replication timing of the two chromosome 1 homologs using a BrdU incorporation assay [see ^57^]. Cultures of cells were pulsed with BrdU for 5.5 hours and mitotic cells harvested, processed for BrdU incorporation and subjected to FISH using a chromosome 1 paint probe plus a BAC probe from within the deleted region (Table S2). As expected, prior to deletion of the TL:1-187 locus, the two chromosome 1 homologs display synchronous replication (see Fig. 6E). In contrast, cells containing heterozygous deletions of either allele resulted in significantly more BrdU incorporation in the deleted chromosome than the non-deleted chromosome, indicating that deletion of either TL1-187 allele results in delayed replication timing in HTD114 cells (Fig. 6E). These results are consistent with the observation that both alleles of TL:1-187 are expressed, and that this locus controls replication timing of human chromosome 1. Therefore, we name this lncRNA gene *ASAR1-187*.

A similar set of RNA-DNA FISH, CRISPR/Cas9 deletion, and BrdU incorporation assays indicated that TL deletions affecting expressed alleles, including both alleles of TL:8-2.7 and single alleles of TL:9-23, TL:9-24, and TL9-30, resulted in delayed replication of their parent chromosomes in *cis*, and we name these genes *ASAR8-2.7*, *ASAR9-23*, *ASAR9-24*, and *ASAR9-30*, respectively (Figures S3 and S4).

### Chromosome structure instability following disruption of TL:1-187

One dramatic phenotype associated with chromosomes with disrupted ASARs is instability of the affected chromosomes, which is characterized by an increase in the rate of chromosomal rearrangements ^7^, an increase in non-disjunction events resulting in cells with anaphase bridges and lagging chromosomes ^8,9^, and an increase in endoreduplication resulting in an increase in the ploidy of the affected cells ^8^. During the characterization of chromosomes with the ASAR deletions described here, we detected extremely late DNA replication [also called “G2 DNA synthesis” ^58,59^], delayed mitotic chromosome condensation, and lagging chromosomes during anaphase. Figure 6 G-J shows an example of an anaphase cell with a lagging chromosome affecting chromosome 1 that is deleted for *ASAR1-187*, and the lagging chromosome also displays delayed mitotic chromosome condensation and extremely late DNA replication with BrdU incorporation that occurred during G2, note that BrdU incorporation was not detected in any other chromosome. Another example of an abnormal anaphase cell with a lagging chromosome 1, also deleted for *ASAR1-187,* is shown in Figure 6K-M, and shows abnormal mitotic condensation in the large heterochromatic region on chromosome 1. Given the prevalence of these abnormal mitotic figures it is not surprising that we also detected numerous rearrangements involving chromosome 1 in cells with deletions of *ASAR1-187* in either CHR1A or CHR1B (Fig. 6N-O).

## Discussion

Chromosome associated lncRNAs have become well established as regulators of chromosome scale replication timing, gene expression and structural stability ^59,60^. The three previously identified ASAR genes display random epigenetically programed AEI, epigenetically programed ASRT, contain a high L1 content, and express RNAs that remain associated with the chromosome territories where they are transcribed ^4-6,9^. In this report, we identified 68 autosomal TL that share all of these “ASAR characteristics”, and genetic disruption of five ASAR candidates resulted in delayed replication timing of human chromosomes in *cis*. These results indicate that ASAR genes are abundant throughout the human genome, are located in regions of the genome that display epigenetically controlled AEI and ASRT, and function to promote proper replication timing and structural stability of each human chromosome.

### Chromosomal loci with variable epigenetic expression and replication timing

In the present study, we used haplotype phased expression and replication timing assays on multiple single-cell derived LCL clones isolated from two unrelated individuals to identify loci that display AEI and ASRT. The clonal and allele-specific analysis of the expression and replication timing profiles for the autosomes revealed a novel genomic behavior at ∼200 loci with differences in AEI and ASRT that are comparable to the differences observed between the active and inactive X chromosomes. These autosomal loci contain both protein coding and noncoding genes that are expressed from single alleles in some clones, expressed from the opposite allele in other clones, expressed from both alleles in other clones, and are not expressed from either allele in yet other clones. The stochastic nature of the AEI at these loci indicates that each allele had acquired one of two states, expressed or silent, and that each allele chose between these two states independently from the other allele. Because this type of AEI is mitotically stable and is not predetermined by parent of origin, we refer to the choice of the on-off state as being random.

Asynchronous replication timing between alleles was originally observed for loci containing olfactory receptor ^61^ and adaptive immune system ^31^ gene arrays, genes that are clearly associated with the exquisite specificity provided by mono-allelic expression. Subsequent studies suggested that all epigenetically programmed mono-allelic genes display asynchronous replication between alleles ^28,31,32,52^. In the present study, we found that each allele can adopt earlier or later replication timing in different clones, which is independent of the other allele. This variability in replication timing results in some clones with dramatic ASRT between alleles, but in other clones the two alleles have synchronous replication, with both alleles adopting either the earlier or later replication timing. Given this wide variability in replication timing we refer to these loci as having <u>V</u>ariable <u>E</u>pigenetic <u>R</u>eplication <u>T</u>iming (VERT). These observations are also consistent with a previous study that identified “megabase sized” regions of <u>A</u>synchronously <u>R</u>eplicating <u>D</u>omains (ARDs) that contain imprinted genes on human autosomes ^1^. The relationship between ARDs and VERTs is currently not clear, but several ARDs overlap with regions described here as having VERT. In addition, the stochastic nature of the allelic expression and replication timing present at VERT loci indicates that each allele acquires expression and replication timing via an epigenetic mechanism that is not dependent on parent of origin, not dependent on the expression or replication timing status of the opposite allele, and the choice of which allele is expressed is not dependent on replication timing. Thus, the Early/Late pattern of asynchronous replication detected in this study is not corelated with which allele is expressed, further supporting the conclusion that asynchronous replication is not a consequence of transcription of only one allele ^4,9,28,32^.

The ASRT of random mono-allelic genes has been proposed to be coordinated with other random mono-allelic genes on the same chromosome ^4,9,28,32,62^. The results described here do not support coordinated ASRT either on the same chromosome nor throughout the genome. Our analysis of the RT profiles from the published mouse pre-B cell data ^53^ shows evidence of VERT, with numerous examples of clone-specific ASRT (see Fig. S2), but the Early/Late RT profiles between the two alleles do not support coordinated ASRT on the same chromosome nor does it support a “mirror image” pattern of ASRT throughout the genome. Instead, we find that each allele can choose between earlier or later replication independently from the other allele, and independently from the other asynchronous loci on the same chromosome.

### Cellular mosaicism is generated by epigenetic AEI of protein coding genes

The variable epigenetic AEI of protein coding genes described here is consistent with an earlier study that also used a clonal analysis and allele-specific expression assay to detect AEI of protein coding genes in human LCLs ^21^. This heterogeneity in expression state at autosomal protein coding genes is thought to introduce cellular mosaicism in what would be otherwise similar cell populations, and has also been referred to as ‘clone-specific’ mono-allelic expression ^63^. Our results are consistent with this interpretation and indicate that expression of each allele is independent of the other allele resulting in bi-allelic as well as silent clones, implying that the mosaicism that is generated is even more extensive than previously believed. The protein coding genes with random epigenetic AEI described here are consistent with previous observations that indicate that random mono-allelic expression of clustered Pcdh genes, which are homophilic cell adhesion molecules, are critical for the generation of cellular individuality in the nervous system^64,65^.

### X chromosome TLs with ASAR characteristics

In this report, we identified 85 TL expressed from the X chromosome, with 66 being expressed exclusively from the active X chromosome and 19 expressed from both the active and inactive X chromosomes. The bi-allelic X chromosome TL reside within regions that are known to escape X inactivation ^43-45^. The hypothesis that the X chromosome contains *cis*-acting loci important for X chromosome inactivation that are separate from the X inactivation center was initially proposed by Drs. Stan Gartler and Arthur Riggs, which they called the “Way Station” model for X inactivation^66^. Subseqeuntly, Dr. Mary Lyon proposed that L1s represent “Booster Elements” that function during the spreading of Xist RNA in *cis* along the X chromosome faciliting the process of inactivation ^67,68^. This notion was supported by the observation that the X chromosome contains ∼27% and autosomes contain ∼13% L1 derived sequences ^69^. In addition, L1s are present at a lower concentration in regions of the X chromosome that escape inactivation, supporting the hypothesis that L1s serve as signals to propagate inactivation along the X chromosome ^69^. Further support for a role of L1s in mono-allelic expression came from the observation that L1s are present at a relatively high local concentration (>18%) near both imprinted and random mono-allelic genes located on autosomes ^37^.

L1 RNA was previously shown to be a major component of C0t1 RNA, which is excluded from heterochromatin and associated with euchromatin ^70,71^. Silencing of C0t1 RNA has been used as a convenient assay for the gene silencing function of *XIST* transgenes when integrated into autosomes ^72,73^. Furthermore, *Xist* mediated silencing of C0t1 RNA expressed from the future inactive X chromosome precedes protein coding gene silencing during early mouse development^74^. We recently used ectopic integration of transgenes and CRISPR/Cas9-mediated chromosome engineering and found that L1 sequences, oriented in the antisense direction, mediate the chromosome-wide effects of *ASAR6* and *ASAR15* ^75^. In addition, oligonucleotides targeting the antisense strand of the one full length L1 within ASAR6 RNA restored normal replication timing to mouse chromosomes expressing an *ASAR6* transgene ^75^. These results provided the first direct evidence that L1 antisense RNA plays a functional role in replication timing of mammalian chromosomes. Taken together, these observations suggest that one role of *XIST* mediated X inactivation involves silencing of ASAR counterparts (i. e. X chromosome TL RNA genes) on the X chromosome, and that regions of the X chromosome that escape inactivation do so by maintaining expression of these L1 rich RNAs that promote the maintenance of euchromatin within the escape regions. Under this scenario the L1 rich TL do not function as “Booster Elements” on the X chromosome, but as targets of XIST-mediated silencing that precedes inactivation of the protein coding genes on the future inactive X chromosome.

In addition to the autosomal loci that display VERT, we detected four regions on the X chromosome, two on the active X and two on the inactive X, that showed allele-specific epigenetic variability in replication timing (see Fig. 3G-J). All four of these regions contain TL with strong AEI, consistent with haplotype 1 representing the expressed and therefore the active X allele. In addition, one of these regions contains two loci that are known to be important for X inactivation in humans. The XACT lncRNA gene is expressed from the active X chromosome and competes with XIST early in human development for the control of X chromosome activity ^48,49^. We note here that the XACT gene is located within a large TL with an annotated vlincRNA [vlinc483; ^36^] that is expressed from the active X chromosome in all six EB3_2 clones (i. e. haplotype 1; Fig. 3J and Table S1). The relationship between the spliced XACT lncRNA product and the larger non-spliced TL is unknown at this time. In addition to XACT, this locus contains DXZ4 and expresses the DANT2 lncRNA ^76^ from the active X chromosome in all six EB3_2 clones (Fig. 3J). DXZ4 is a macrosatellite repeat that lies at the boundary of a “superdomain” on the inactive X chromosome, and the inactive X allele extensively binds CTCF ^50^. Deletion of DXZ4 leads to the disappearance of “superdomains”, changes in compartmentalization patterns, and changes in the distribution of chromatin marks on the inactive X chromosome ^51^. The presence of VERT loci on the active and inactive X chromosomes suggests that the epigenetic program described here is not limited to autosomes but also functions on the X chromosomes in female cells. Whether or not VERT is present on the single X chromosome in male cells is currently not known.

### Protein coding genes that display epigenetically programmed AEI

In the immune, olfactory, and central nervous systems, cellular individuality is generated by expression of diverse receptor-type molecules, and involves random mono-allelic expression resulting in specific functional properties. To ensure that individual B or T cells express single Ig or TCR genes, respectively, mono-allelic expression functions to limit the number of expressed genes to one per cell ^31^. At the molecular level, the mechanism of stochastic mono-allelic expression can help generate cellular individuality and provides potential functional variation among the individual cells of a complex system ^23,63,64^. The protein coding genes that display epigenetically controlled AEI described here are primarily involved in homophilic cell adhesion, suggesting that B cells may also use mono-allelic expression of cell adhesion molecules in addition to the Ig genes during the adaptive immune response.

## Conclusions

Genomes are defined by their sequence. However, the linear arrangement of nucleotides along chromosomes is only their most basic feature. Within this linear arrangement of nucleotides exists three different types of *cis*-acting loci known to be essential for normal function of every eukaryotic chromosome; origins of replication, centromeres, and telomeres are present on all linear chromosomes functioning to promote proper replication, segregation and stability of each chromosome. The identification of ASARs present on all human chromosomes and essential for timely replication, condensation and genetic stability supports a fourth type of essential *cis*-acting chromosomal locus, which we call “Inactivation/Stability Centers (I/SCs)”. I/SCs likely are present in all mammals and may pre-date the evolution of *XIST*, which is a critical component of the X inactivation center present in eutherian mammals but absent in metatherian mammals ^77^. I/SCs are complex loci that contain protein coding genes that are important for cell-cell adhesion and cellular individuality, contain multiple non-coding ASAR genes that are expressed in different tissues and are responsible for proper replication timing and structural stability of each chromosome.

## Abbreviations

ASAR: Asynchronous Replication and Autosomal RNA
AEI: Allelic Expression Imbalance
RT: Replication Timing
ASRT: Asynchronous Replication Timing
TL: Transcribed Loci
eQTL: expression Quantitative Trait Loci
rtQTL: replication timing Quantitative Trait Loci
vlincRNA: very long intergenic non-coding RNA
ENCODE: Encyclopedia of DNA Elements
LCL: lymphoblastoid cell line
BAC: Bacterial Artificial Chromosome
VERT: Variable Epigenetic Replication Timing
ARD: Asynchronously Replicating Domain

## Acknowledgments

M.J.T. was supported by NIH NIGMS (R01GM114162 and R01GM130703).

M.B.H was supported by NIH NCI 4K00CA245677-03

---

**Figure S1.**
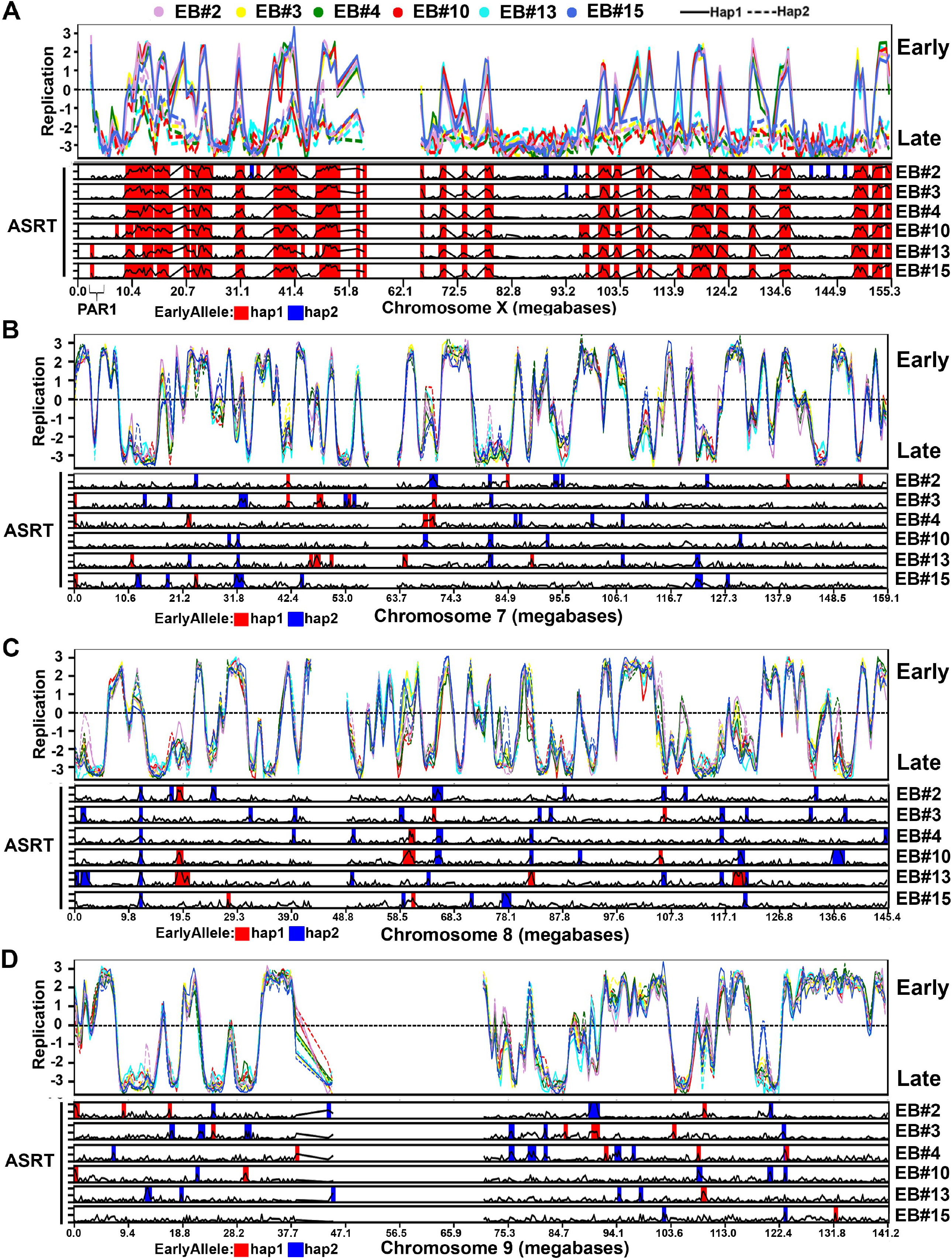
Asynchronous Replication Timing on human chromosomes. Chromosome Early/Late RT profiles from the 6 EB3_2 clones highlighting regions with SD >1 from the asynchronous replication timing (ASRT) analysis on individual clones. Each clone was color coded as shown, with haplotype 1 shown as a solid line, and haplotype 2 shown as a dotted line for each clone. The left axis shows the RT profiles, with positive numbers representing early replication and negative numbers representing late replication. Early replicating loci with SD >1 are highlighted in red for haplotype 1 and in blue for haplotype 2. The position on each chromosome is shown in magabases. Chromosome profiles are shown for the X chromosome (A), chromosome 7 (B), chromosome 8 (C) and chromosome 9 (D).

**Figure S2.**
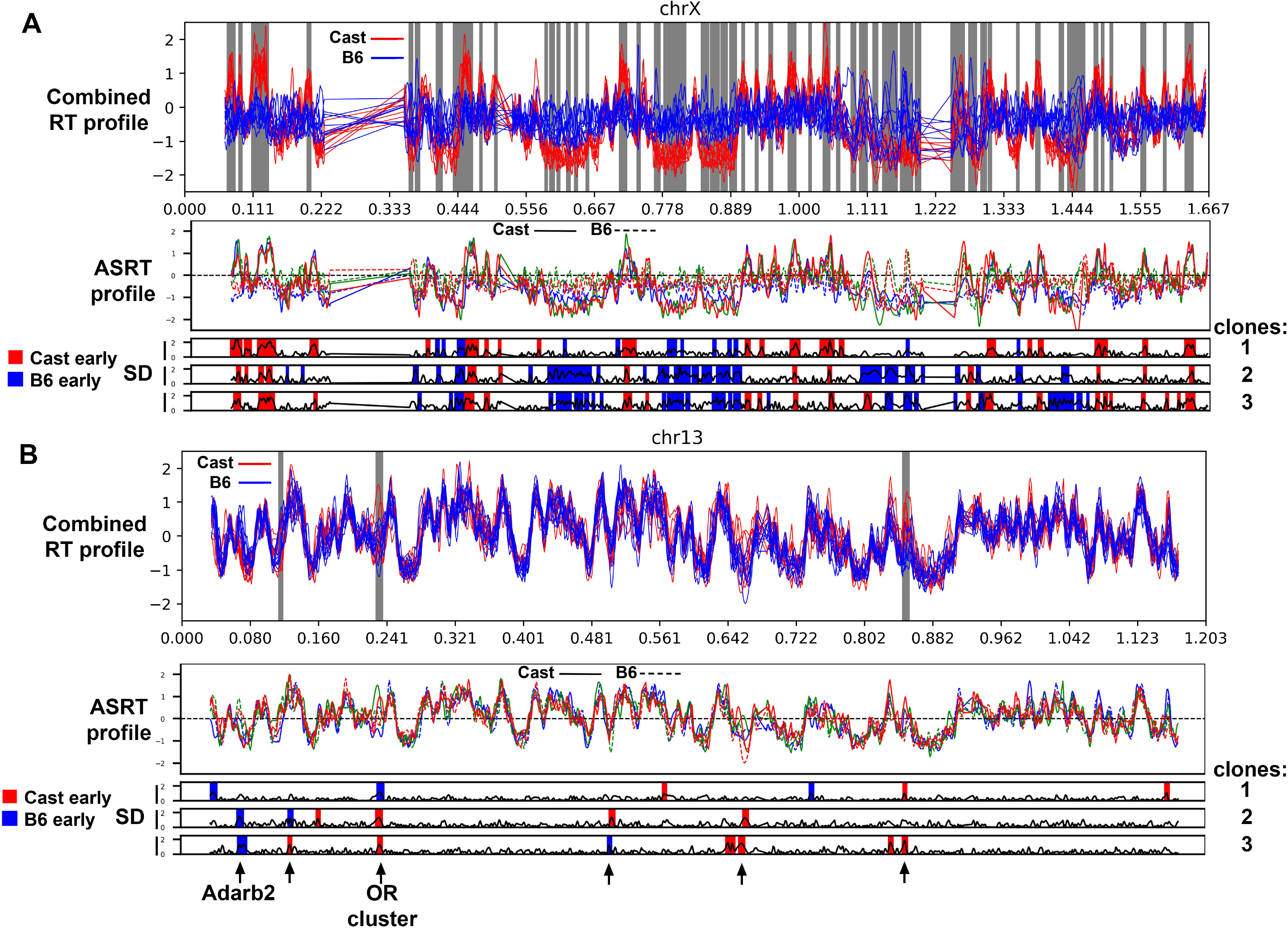
Asynchronous Replication Timing on mouse chromosomes. Chromosome Early/Late RT profiles from 3 pre-B cell clones [from ^53^] showing the “combined RT profile” analysis (Top panels) and the asynchronous replication timing profile (ASRT) analysis on individual clones (Middle panels), regions with SD >1 from the ASRT analysis is shown below each panel. For the “combined RT profile” analysis the C57BL6 (B6) allele is blue and the Castaneous (Cast) allele is red, and the regions with SD >1 are highlighted in gray. For the ASRT analysis, each clone was color coded, with the C57BL6 allele as a solid line and the Castaneous allele as a dotted line. The left axis shows the RT profiles, with positive numbers representing early replication and negative numbers representing late replication. From the ASRT analysis early replicating loci with SD >1 are highlighted in red for the Castaneous allele and in blue for C57BL6 allele. The position on each chromosome is shown in magabases. Chromosome profiles are shown for the X chromosome (A) and chromosome 13 (B). The arrows mark 6 loci on chromosome 13 that show ASRT in multiple clones. We detected ASRT at loci that are known to be subject to random monoallelic expression, e. g. an olfactory receptor cluster (OR cluster) and Adarb2 (see Fig. 5A).

**Figure S3.**
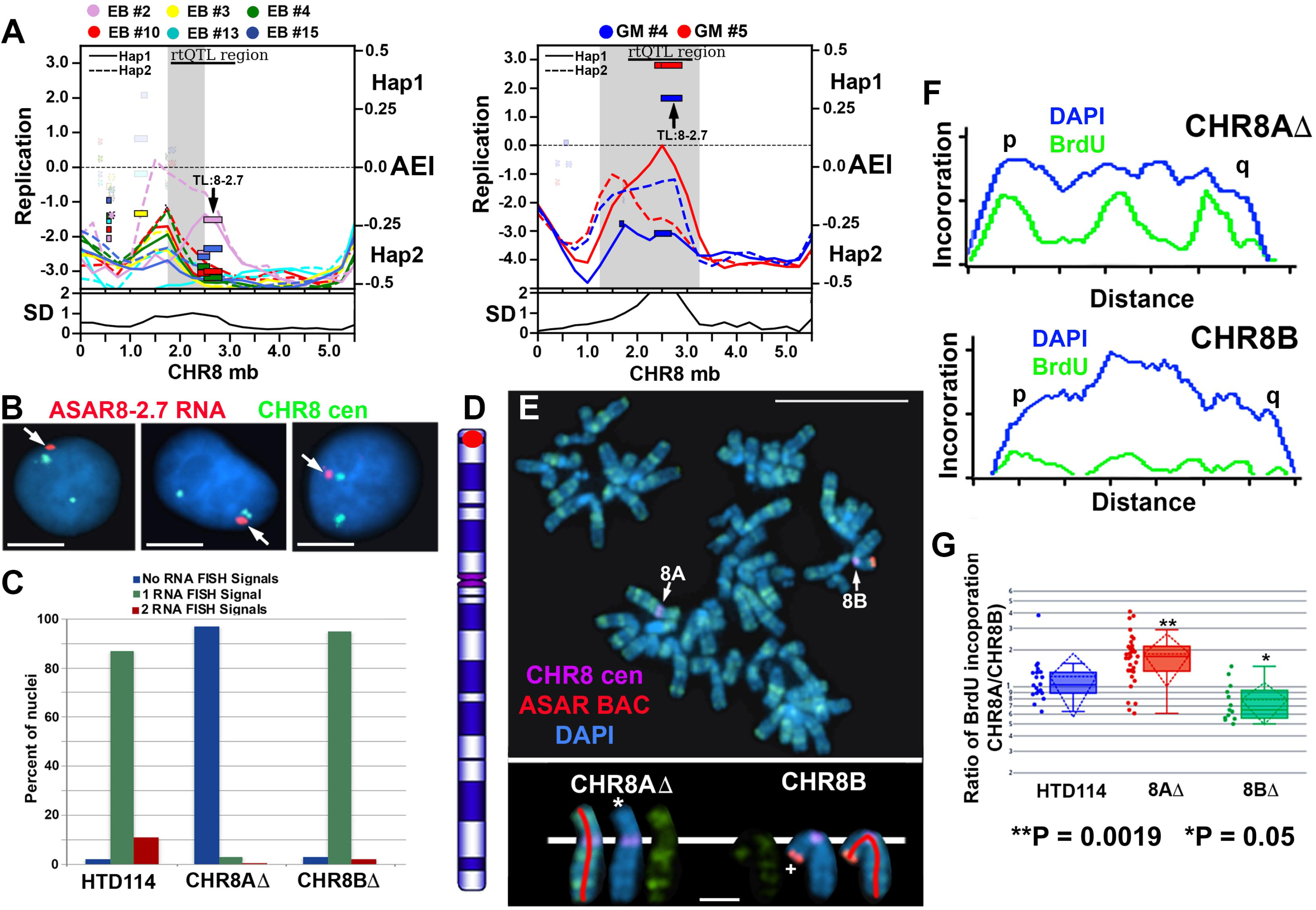
Delayed replication of chromosome 8 following disruption of TL:8-2.7. A) AEI and ASRT at the TL:8-2.7 locus in the EB3_2 (left panel) and GM12878 (right panel) clones. The left axis shows the replication timing (Early/Late) profiles, and the standard deviation (SD) across the locus is shown below each panel. The right axis shows the AEI of haplotype 1 (Hap1) and haplotype 2 (Hap2). The smooth rectangles represent TL, with TL:8-2.7 marked with arrows. The stippled rectangles represent protein coding genes. The opaque rectangles show AEI (FDR-BH alpha <=0.01), and the transparent rectangles show bi-allelic expression of protein coding and non-coding transcripts. B) Examples of cells analyzed using RNA-DNA FISH using at TL:8-2.7 probe (red; see Table S2) to detect RNA and a chromosome 8 centromeric (green; CHR8 cen) probe to detect DNA. Arrowheads mark the sites of TL:8-2.7 RNA FISH hybridization. C) Quantitation of the RNA FISH hybridization signals for TL:8-2.7 before and after allele-specific deletion of the locus in the HTD114 cell line. D) BrdU incorporation in HTD114 cells containing a heterozygous deletion of the TL:8-2.7 locus. Cells were processed for BrdU incorporation and processed for DNA FISH using a chromosome 8 centromeric probe (purple) plus a Fosmid from within the deleted region (red; see Table S2). The TL:8-2.7 Fosmid was used to identify the chromosome 8 with the deletion. E) The pixel intensity profiles for both DAPI (blue) and BrdU (green) staining for the two chromosome 8s from panel D are shown. The short (p) and long (q) arms of chromosome 8 are indicated. Note the BrdU incorporation is greater in the deleted chromosome (CHR8AΔ) than in the non-deleted chromosome (CHR8B). Quantification of BrdU incorporation in multiple cells with heterozygous deletions indicated that deletion of either allele resulted in delayed replication of chromosome 8 (8AΔ or 8BΔ). The P values form the Kruskal-Wallis test ^78^ are shown.

**Figure S4.**
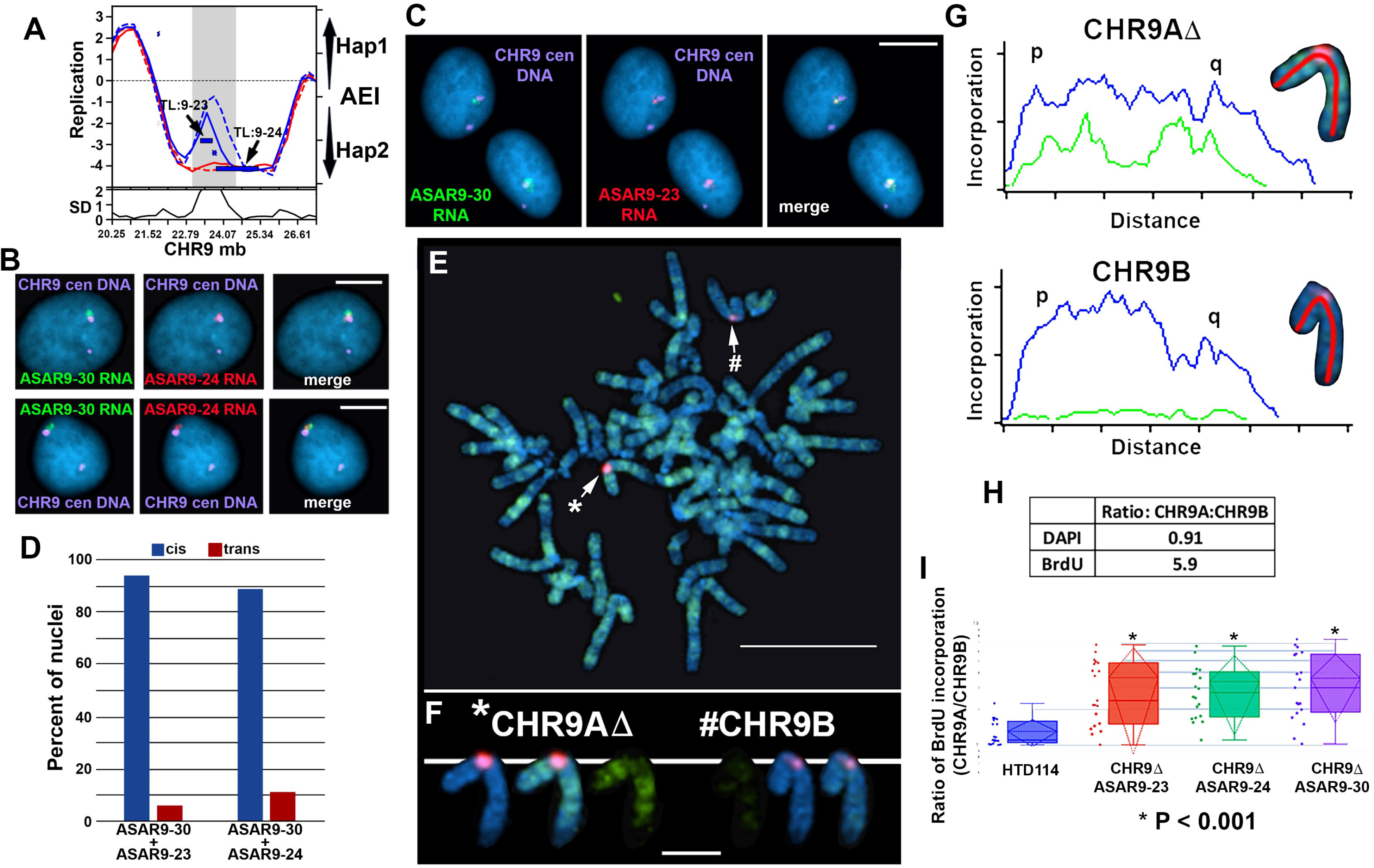
Delayed replication of chromosome 9 following disruption of TL:9-23, TL:9-24 and TL:9-30. A) AEI and ASRT at the TL:9-23 and TL:9-24 locus in the GM12878 clones. The left axis shows the replication timing (Early/Late) profiles, and the standard deviation (SD) across the locus is shown below. The right axis shows the AEI of haplotype 1 (Hap1) and haplotype 2 (Hap2). The smooth rectangles represent TL, with TL:9-23 and TL:9-24 are marked with arrows. The stippled rectangles represent protein coding genes. The opaque rectangles show AEI (FDR-BH alpha <=0.01), and the transparent rectangles show bi-allelic expression of protein coding and non-coding transcripts. B) Examples of cells analyzed using RNA-DNA FISH using at TL:9-24 (ASAR9-24) probe (red) and TL:9-30 (ASAR9-30) probe (green) to detect RNA and a chromosome 9 centromeric (purple; CHR9 cen) probe to detect DNA. Arrowheads mark the sites of TL:8-2.7 RNA FISH hybridization. C) Examples of cells analyzed using RNA-DNA FISH using at TL:9-23 (ASAR9-23) probe (red) and TL:9-30 (ASAR9-30) probe (green) to detect RNA and a chromosome 9 centromeric (purple; CHR9 cen) probe to detect DNA. Arrowheads mark the sites of TL:8-2.7 RNA FISH hybridization. D) Expression of TL:9-23 (ASAR9-23), TL:9-24 (ASAR9-24), and TL:9-30 (ASAR9-30) occurs from the same chromosome 9 homolog in HTD114 cells. RNA FISH hybridization signals were detected in the same nuclei and scored for expression from the same chromosome 9 homolog (i. e. in *cis*) or opposite homologs (i. e. in *trans*). E) BrdU incorporation in HTD114 cells containing a heterozygous deletion of the TL:9-23 locus. Cells were processed for BrdU incorporation and processed for DNA FISH using a chromosome 9 centromeric probe (red). Note that the centromeric probe detects a polymorphism in the size of the FISH signal, and the chromosome with the larger signal is referred to as CHR9A (*) which is the expressed allele for all three TL. The chromosome with the smaller centromeric signal is referred to as CHR9B (#) and is silent for all 3 TL. E) The pixel intensity profiles for both DAPI (blue) and BrdU (green) staining for the two chromosome 9s from panel F are shown. The short (p) and long (q) arms of chromosome 9 are indicated. Note the BrdU incorporation is greater in the deleted chromosome (CHR9AΔ) than in the non-deleted chromosome (CHR9B). Quantification of BrdU incorporation in multiple cells with heterozygous deletions indicated that deletion of either allele resulted in delayed replication of chromosome 9 (CHR9ΔASAR9-23, CHR9ΔASAR9-24, CHR9ΔASAR9-30). The P values are from the Kruskal-Wallis test ^78^.

## STAR Methods

### Cell culture

GM12878 cells were obtained from ATCC and EB3_2 cells were from ^46^. LCLs were grown in RPMI 1640 (Life Technologies) supplemented with 15% fetal bovine serum (Hyclone). Single cell clones were isolated by plating individual cells in 96 well dishes, and were expanded for >25 population doublings. Primary blood lymphocytes were isolated after venipuncture into a Vacutainer CPT (Becton Dickinson, Franklin Lakes, NJ) per the manufacturer’s recommendations and grown in 5 mL RPMI 1640 (Life Technologies) supplemented with 10% fetal bovine serum (Hyclone) and 1% phytohemagglutinin (Life Technologies). HTD114 cells are a human *APRT* deficient cell line derived from HT1080 cells ^79^, and were grown in DMEM (Gibco) supplemented with 10% fetal bovine serum (Hyclone). All cells were grown in a humidified incubator at 37°C in a 5% carbon dioxide atmosphere.

### RNA-seq

Nuclei were isolated by centrifugation for 0.5 minutes from GM12878 and EB3_2 clones following lysis in 0.5% NP40, 140 mM NaCl, 10 mM Tris-HCl (pH 7.4), and 1.5 mM MgCl_2_. Nuclear RNA was isolated using Trizol reagent using the manufacturer’s instructions, followed by DNase treatment to remove possible genomic DNA contamination. Briefly, ribosomal RNAs were removed using the Ribo-Zero kit (Illumina), RNA was fragmented into 250-300bp fragments, and cDNA libraries were prepared using the Directional RNA Library Prep Kit (NEB). Paired end sequencing was done on a NovaSeq 6000 at the OHSU MPSSR core facility. Sequences were aligned to the human genome (hg19) using the STAR aligner ^80^ with default settings. Duplicate reads and reads with map quality below 30 were removed with SAMtools ^81^.

### Early/Late Repli-seq

Early/Late Repli-seq librares were generated as previously described ^82^. In brief, cells were pulse labeled with BrdU, then sorted by flow cytometry according to DNA content into an early and late S-phase bins. Nascent DNA is then immunoprecipitated from both early and late fractions, and standard Illumina sequencing was performed.

### Identification of Transcribed Loci (TLs)

Nuclear-enriched, strand-specific, ribo-minus, total-RNA sequencing libraries were first aligned to the hg19 reference genome using default parameters with STAR ^80^. Next SAMtools ^81^ was used to remove duplicate and low quality (<= MAPQ20) reads. TLs were defined using a strategy of sequential merging of strand-specific, contiguous intergenic reads, as described previously ^34-36^. For each cell line, TLs were defined after combining aligned reads from all derivative subclones, and considered expressed in each subclone if a minimum of 20 informative reads (i.e. covering a heterozygous SNP locus) were detected.

### Allelic expression imbalance analysis of TLs and coding genes in GM12878 and EB3_2

AEI was defined as the fraction of strand-concordant, allele-specific reads to the total number of informative reads. Thus, perfect allelic balance yields a value of 0.5, and perfect allelic bias towards one haplotype yields a value of 1. For convenience, allelic bias from Haplotype 1 is plotted as positive values, allelic bias from Haplotype 2 is plotted as negative values, and AEI is transformed to percentages in the figures.

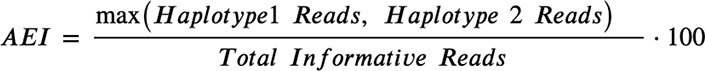

A fully haplotype resolved reference genotype for GM12878 was obtained from the Illumina Platinum Genome project ^41^, and a fully haplotype resolved reference genotype for EB3_2 was previously described ^46^.

TLs and genes with strong AEI were defined as the union of outliers identified by a parametric and non-parametric statistical outlier identification method: 1) Two-sided Binomial-test p-value<=0.001 and FDR Benjamini-Hochberg adjusted q-value<=0.01; and 2) To address the increase of variance with higher expressed genes, linear regression was used to model the relationship between AEI and expression level for all annotated expressed genes and TLs. Outliers were defined as expressed TLs or genes displaying AEI greater than the predicted transcriptome-wide average value plus 2.5 times the standard deviation of AEI for a given expression level.

### Allele-specific DNA replication-timing analysis

∼10x whole genome sequencing was performed on the Early and Late fractions of each Repli-seq sample and reads aligned using BWA MEM ^83^. Utilizing fully haplotype-resolved reference genotypes, allele-specific Repli-seq reads were enumerated for each allele of the Early and Late sequencing fractions. After quantile normalization, the Log2(Early/Late) ratio was calculated for non-overlapping genomic windows of 250kb to create a genome-wide replication timing profile for each haplotype. To identify regions with epigenetic variation between alleles, the standard deviation of replication timing among alleles, SD_alleles,_ was calculated for each genomic window. Windows were considered outliers if displaying an SD_alleles_ value 2.5 standard deviations over the median SD_alleles_ value of all genomic windows measured.

### PCR and expression analysis

Genomic DNA and total RNA was isolated from tissue culture cells using TRIZOL Reagent (Invitrogen). cDNA was prepared using the SuperScript^TM^ III First-Strand Synthesis System (Invitrogen). Reverse transcriptase reactions were performed in the presence or absence of reverse transcriptase on 5µg of total RNA. PCR (genomic and RT-PCR) was performed in a 25-50µL volume using 50-100ng of genomic DNA or 1-2 µL of cDNA (50-100ng of input RNA equivalent), 1x Standard Taq Buffer (New England Biolabs, Inc.), 200µM each deoxynucleotide triphosphates, 0.2µM of each primer, and 3 units of Taq DNA Polymerase (New England Biolabs, Inc.) under the following reaction conditions: 95°C for 2 minutes, followed by 30-40 cycles of 95°C for 30 seconds, 55-62°C for 45 seconds, and 72°C for 1 minute, with a final extension time of 10 minutes at 72°C. PCR products were separated on 1% agarose gels, stained with ethidium bromide, and photographed under ultraviolet light illumination. Sequencing of PCR products was carried out at the Vollum Institute DNA Sequencing Core facility.

### DNA FISH

Mitotic chromosome spreads were prepared as described previously ^84^. After RNase (100µg/ml) treatment for 1h at 37^°^C, slides were washed in 2XSSC and dehydrated in an ethanol series and allowed to air dry. Chromosomal DNA on the slides was denatured at 75^°^C for 3 minutes in 70% formamide/2XSSC, followed by dehydration in an ice-cold ethanol series and allowed to air dry. BAC and Fosmid DNAs were labeled using nick translation (Vysis, Abbott Laboratories) with Spectrum Orange-dUTP, Spectrum Aqua-dUTP or Spectrum Green-dUTP (Vysis). Final probe concentrations varied from 40-60 ng/µl. Centromeric probe cocktails (Vysis) and/or whole chromosome paint probes (Metasystems) plus BAC or Fosmid DNAs were denatured at 75^°^C for 10 minutes and prehybridized at 37^°^C for 10 minutes. Probes were applied to denatured slides and incubated overnight at 37^°^C. Post-hybridization washes consisted of one 3-minute wash in 50% formamide/2XSSC at 40^°^C followed by one 2-minute rinse in PN (0.1M Na_2_HPO_4_, pH 8.0/2.5% Nonidet NP-40) buffer at RT. Coverslips were mounted with Prolong Gold antifade plus DAPI (Invitrogen) and viewed under UV fluorescence (Olympus).

### RNA-DNA FISH

Cells were plated on Poly-L-Lysine coated (Millipore Singa) glass microscope slides at ∼50% confluence and incubated for 4 hours in complete media in a 37°C humidified CO_2_ incubator. Slides were rinsed 1X with sterile RNase free PBS. Cell Extraction was carried out using ice cold solutions as follows: Slides were incubated for 30 seconds in CSK buffer (100mM NaCl/300mM sucrose/3mM MgCl_2_/10mM PIPES, pH 6.8), 10 minutes in CSK buffer/0.1% Triton X-100, followed by 30 seconds in CSK buffer. Cells were then fixed in 4% paraformaldehyde in PBS for 10 minutes and stored in 70% EtOH at −20°C until use. Just prior to RNA FISH, slides were dehydrated through an EtOH series and allowed to air dry. Denatured probes were prehybridized at 37°C for 10 min, applied to non-denatured slides and hybridized at 37°C for 14-16 hours. Post-hybridization washes consisted of one 3-minute wash in 50% formamide/2XSSC at 40^°^C followed by one 2-minute rinse in 2XSSC/0.1% TX-100 for 1 minute at RT. Slides were then fixed in 4% paraformaldehyde in PBS for 5 minutes at RT, and briefly rinsed in 2XSSC/0.1% TX-100 at RT. Coverslips were mounted with Prolong Gold antifade plus DAPI (Invitrogen) and slides were viewed under UV fluorescence (Olympus). Z-stack images were generated using a Cytovision workstation. After capturing RNA FISH signals, the coverslips were removed, the slides were dehydrated in an ethanol series, and then processed for DNA FISH, beginning with the RNase treatment step, as described above.

### Chromosome replication timing assay

The BrdU replication timing assay was performed as described previously on exponentially dividing cultures and asynchronously growing cells ^85^. Mitotic chromosome spreads were prepared and DNA FISH was performed as described above. The incorporated BrdU was then detected using a FITC-labeled anti-BrdU antibody (Roche). Coverslips were mounted with Prolong Gold antifade plus DAPI (Invitrogen), and viewed under UV fluorescence. All images were captured with an Olympus BX Fluorescent Microscope using a 100X objective, automatic filter-wheel and Cytovision workstation. Individual chromosomes were identified with either chromosome-specific paints, centromeric probes, BACs or by inverted DAPI staining. Utilizing the Cytovision workstation, each chromosome was isolated from the metaphase spread and a line drawn along the middle of the entire length of the chromosome. The Cytovision software was used to calculate the pixel area and intensity along each chromosome for each fluorochrome occupied by the DAPI and BrdU (FITC) signals. The total amount of fluorescent signal in each chromosome was calculated by multiplying the average pixel intensity by the area occupied by those pixels. The BrdU incorporation into human chromosomes containing CRISPR/Cas9 modifications was calculated by dividing the total incorporation into the chromosome with the deleted chromosome divided by the BrdU incorporation into the non-deleted chromosome within the same cell. Boxplots were generated from data collected from 8-12 cells per clone or treatment group. Differences in measurements were tested across categorical groupings by using the Kruskal-Wallis test ^78^ and listed as P-values for the corresponding plots.

### CRISPR/Cas9 engineering

Using Lipofectamine 2000, according to the manufacturer’s recommendations, we co-transfected HTD114 cells with plasmids encoding GFP, sgRNAs and Cas9 endonuclease (Origene). Each plasmid encoded sgRNA was designed to bind at the indicated locations (Table S4). 48h after transfection, cells were plated at clonal density and allowed to expand for 2-3 weeks. The presence of deletions was identified by PCR using the primers described in Table S4. The single cell colonies that grew were analyzed for heterozygous deletions by PCR. We used retention of heterozygous SNPs (see Table S4) to identify the disrupted allele, and homozygosity at these SNPs confirmed that cell clones were homogenous.

## RESOURCE AVAILABILITY

Further information and requests for resources and reagents should be directed to and will be fulfilled by the lead contact, Mathew Thayer

### Materials availability statement

All reagents generated in this study will be made freely available upon request.

## References

1. Mukhopadhyay, R. et al. Allele-specific genome-wide profiling in human primary erythroblasts reveal replication program organization. PLoS Genet 10, e1004319 (2014).

2. Dileep, V. & Gilbert, D.M. Single-cell replication profiling to measure stochastic variation in mammalian replication timing. Nat Commun 9, 427 (2018).

3. Rivera-Mulia, J.C. et al. Allele-specific control of replication timing and genome organization during development. Genome Res 28, 800–811 (2018).

4. Donley, N., Smith, L. & Thayer, M.J. ASAR15, A cis-Acting Locus that Controls Chromosome-Wide Replication Timing and Stability of Human Chromosome 15. PLoS Genet 11, e1004923 (2015).

5. Heskett, M.B., Smith, L.G., Spellman, P. & Thayer, M.J. Reciprocal monoallelic expression of ASAR lncRNA genes controls replication timing of human chromosome 6. Rna 26, 724–738 (2020).

6. Stoffregen, E.P., Donley, N., Stauffer, D., Smith, L. & Thayer, M.J. An autosomal locus that controls chromosome-wide replication timing and mono-allelic expression. Hum Mol Genet 20, 2366–2378 (2011).

7. Breger, K.S., Smith, L. & Thayer, M.J. Engineering translocations with delayed replication: evidence for cis control of chromosome replication timing. Hum Mol Genet 14, 2813–27 (2005).

8. Chang, B.H., Smith, L., Huang, J. & Thayer, M. Chromosomes with delayed replication timing lead to checkpoint activation, delayed recruitment of Aurora B and chromosome instability. Oncogene 26, 1852–61 (2007).

9. Donley, N., Stoffregen, E.P., Smith, L., Montagna, C. & Thayer, M.J. Asynchronous Replication, Mono-Allelic Expression, and Long Range Cis-Effects of ASAR6. PLoS Genet 9, e1003423 (2013).

10. Platt, E.J., Smith, L. & Thayer, M.J. L1 retrotransposon antisense RNA within ASAR lncRNAs controls chromosome-wide replication timing. J Cell Biol 217, 541–553 (2018).

11. Bryois, J. et al. Cis and trans effects of human genomic variants on gene expression. PLoS Genet 10, e1004461 (2014).

12. Petretto, E. et al. Heritability and tissue specificity of expression quantitative trait loci. PLoS Genet 2, e172 (2006).

13. Alexander, M.K. et al. Differences between homologous alleles of olfactory receptor genes require the Polycomb Group protein Eed. J Cell Biol 179, 269–76 (2007).

14. Gendrel, A.V. et al. Developmental dynamics and disease potential of random monoallelic gene expression. Dev Cell 28, 366–80 (2014).

15. Li, S.M. et al. Transcriptome-wide survey of mouse CNS-derived cells reveals monoallelic expression within novel gene families. PLoS One 7, e31751 (2012).

16. Lin, M. et al. Allele-biased expression in differentiating human neurons: implications for neuropsychiatric disorders. PLoS One 7, e44017 (2012).

17. Gendrel, A.V., Marion-Poll, L., Katoh, K. & Heard, E. Random monoallelic expression of genes on autosomes: Parallels with X-chromosome inactivation. Semin Cell Dev Biol (2016).

18. Bartolomei, M.S. Genomic imprinting: employing and avoiding epigenetic processes. Genes Dev 23, 2124–33 (2009).

19. Tucci, V., Isles, A.R., Kelsey, G. & Ferguson-Smith, A.C. Genomic Imprinting and Physiological Processes in Mammals. Cell 176, 952–965 (2019).

20. Chess, A. Mechanisms and consequences of widespread random monoallelic expression. Nat Rev Genet 13, 421–8 (2012).

21. Gimelbrant, A., Hutchinson, J.N., Thompson, B.R. & Chess, A. Widespread monoallelic expression on human autosomes. Science 318, 1136–40 (2007).

22. Reinius, B. & Sandberg, R. Random monoallelic expression of autosomal genes: stochastic transcription and allele-level regulation. Nat Rev Genet 16, 653–64 (2015).

23. Savova, V. et al. Genes with monoallelic expression contribute disproportionately to genetic diversity in humans. Nat Genet 48, 231–237 (2016).

24. Heskett, M.B., Spellman, P.T. & Thayer, M.J. Differential Allelic Expression among Long Non-Coding RNAs. Noncoding RNA 7 (2021).

25. Ding, Q. et al. The genetic architecture of DNA replication timing in human pluripotent stem cells. Nat Commun 12, 6746 (2021).

26. Koren, A. et al. Genetic variation in human DNA replication timing. Cell 159, 1015–1026 (2014).

27. Zhao, P.A., Sasaki, T. & Gilbert, D.M. High-resolution Repli-Seq defines the temporal choreography of initiation, elongation and termination of replication in mammalian cells. Genome Biol 21, 76 (2020).

28. Ensminger, A.W. & Chess, A. Coordinated replication timing of monoallelically expressed genes along human autosomes. Hum Mol Genet 13, 651–8 (2004).

29. Goldmit, M. & Bergman, Y. Monoallelic gene expression: a repertoire of recurrent themes. Immunol Rev 200, 197–214 (2004).

30. Gribnau, J., Hochedlinger, K., Hata, K., Li, E. & Jaenisch, R. Asynchronous replication timing of imprinted loci is independent of DNA methylation, but consistent with differential subnuclear localization. Genes Dev 17, 759–73 (2003).

31. Mostoslavsky, R. et al. Asynchronous replication and allelic exclusion in the immune system. Nature 414, 221–5 (2001).

32. Singh, N. et al. Coordination of the random asynchronous replication of autosomal loci. Nat Genet 33, 339–41 (2003).

33. St Laurent, G., Savva, Y.A. & Kapranov, P. Dark matter RNA: an intelligent scaffold for the dynamic regulation of the nuclear information landscape. Front Genet 3, 57 (2012).

34. Caron, M. et al. Very long intergenic non-coding RNA transcripts and expression profiles are associated to specific childhood acute lymphoblastic leukemia subtypes. PLoS One 13, e0207250 (2018).

35. St Laurent, G., et al. VlincRNAs controlled by retroviral elements are a hallmark of pluripotency and cancer. Genome Biol 14, R73 (2013).

36. St Laurent, G., et al. Functional annotation of the vlinc class of non-coding RNAs using systems biology approach. Nucleic Acids Res 44, 3233–52 (2016).

37. Allen, E. et al. High concentrations of long interspersed nuclear element sequence distinguish monoallelically expressed genes. Proc Natl Acad Sci U S A 100, 9940–5 (2003).

38. Breger, K.S., Smith, L., Turker, M.S. & Thayer, M.J. Ionizing radiation induces frequent translocations with delayed replication and condensation. Cancer Research 64, 8231–8238 (2004).

39. Moore, J.E. et al. Expanded encyclopaedias of DNA elements in the human and mouse genomes. Nature 583, 699–710 (2020).

40. St Laurent, G., et al. VlincRNAs controlled by retroviral elements are a hallmark of pluripotency and cancer. Genome Biology 14, R73 (2013).

41. Eberle, M.A. et al. A reference data set of 5.4 million phased human variants validated by genetic inheritance from sequencing a three-generation 17-member pedigree. Genome Res 27, 157–164 (2017).

42. Shvetsova, E. et al. Skewed X-inactivation is common in the general female population. Eur J Hum Genet 27, 455–465 (2019).

43. Tukiainen, T. et al. Landscape of X chromosome inactivation across human tissues. Nature 550, 244–248 (2017).

44. Posynick, B.J. & Brown, C.J. Escape From X-Chromosome Inactivation: An Evolutionary Perspective. Front Cell Dev Biol 7, 241 (2019).

45. Navarro-Cobos, M.J., Balaton, B.P. & Brown, C.J. Genes that escape from X-chromosome inactivation: Potential contributors to Klinefelter syndrome. Am J Med Genet C Semin Med Genet 184, 226–238 (2020).

46. Lajugie, J. et al. Complete genome phasing of family quartet by combination of genetic, physical and population-based phasing analysis. PLoS One 8, e64571 (2013).

47. Wardemann, H. et al. Predominant autoantibody production by early human B cell precursors. Science 301, 1374–7 (2003).

48. Vallot, C. et al. XACT, a long noncoding transcript coating the active X chromosome in human pluripotent cells. Nat Genet 45, 239–41 (2013).

49. Vallot, C. et al. XACT Noncoding RNA Competes with XIST in the Control of X Chromosome Activity during Human Early Development. Cell Stem Cell 20, 102–111 (2017).

50. Chadwick, B.P. DXZ4 chromatin adopts an opposing conformation to that of the surrounding chromosome and acquires a novel inactive X-specific role involving CTCF and antisense transcripts. Genome Res 18, 1259–69 (2008).

51. Darrow, E.M. et al. Deletion of DXZ4 on the human inactive X chromosome alters higher-order genome architecture. Proc Natl Acad Sci U S A 113, E4504–12 (2016).

52. Schlesinger, S., Selig, S., Bergman, Y. & Cedar, H. Allelic inactivation of rDNA loci. Genes Dev 23, 2437–47 (2009).

53. Blumenfeld, B. et al. Chromosomal coordination and differential structure of asynchronous replicating regions. Nat Commun 12, 1035 (2021).

54. Bergman, Y., Simon, I. & Cedar, H. Asynchronous Replication Timing: A Mechanism for Monoallelic Choice During Development. Front Cell Dev Biol 9, 737681 (2021).

55. Koren, A. & McCarroll, S.A. Random replication of the inactive X chromosome. Genome Res 24, 64–9 (2014).

56. Raudvere, U. et al. g:Profiler: a web server for functional enrichment analysis and conversions of gene lists (2019 update). Nucleic Acids Research 47, W191–W198 (2019).

57. Smith, L. & Thayer, M. Chromosome replicating timing combined with fluorescent in situ hybridization. J Vis Exp 10, e4400 (2012).

58. zur Hausen, H. Chromosomal changes of similar nature in seven established cell lines derived from the peripheral blood of patients with leukemia. J Natl Cancer Inst 38, 683–96 (1967).

59. Thayer, M.J. Mammalian chromosomes contain cis-acting elements that control replication timing, mitotic condensation, and stability of entire chromosomes. Bioessays 34, 760–70 (2012).

60. Galupa, R. & Heard, E. X-Chromosome Inactivation: A Crossroads Between Chromosome Architecture and Gene Regulation. Annu Rev Genet 52, 535–566 (2018).

61. Chess, A., Simon, I., Cedar, H. & Axel, R. Allelic inactivation regulates olfactory receptor gene expression. Cell 78, 823–34 (1994).

62. Masika, H. et al. Programming asynchronous replication in stem cells. Nat Struct Mol Biol 24, 1132–1138 (2017).

63. Savova, V., Vigneau, S. & Gimelbrant, A.A. Autosomal monoallelic expression: genetics of epigenetic diversity? Curr Opin Genet Dev 23, 642–8 (2013).

64. Yagi, T. Genetic basis of neuronal individuality in the mammalian brain. J Neurogenet 27, 97–105 (2013).

65. Mountoufaris, G. et al. Multicluster Pcdh diversity is required for mouse olfactory neural circuit assembly. Science 356, 411–414 (2017).

66. Gartler, S.M. & Riggs, A.D. Mammalian X-chromosome inactivation. Annu Rev Genet 17, 155–90 (1983).

67. Lyon, M.F. X-chromosome inactivation: a repeat hypothesis. Cytogenet Cell Genet 80, 133–7 (1998).

68. Lyon, M.F. The Lyon and the LINE hypothesis. Semin Cell Dev Biol 14, 313–8 (2003).

69. Bailey, J.A., Carrel, L., Chakravarti, A. & Eichler, E.E. Molecular evidence for a relationship between LINE-1 elements and X chromosome inactivation: the Lyon repeat hypothesis. Proc Natl Acad Sci U S A 97, 6634–9 (2000).

70. Creamer, K.M., Kolpa, H.J. & Lawrence, J.B. Nascent RNA scaffolds contribute to chromosome territory architecture and counter chromatin compaction. Mol Cell (2021).

71. Hall, L.L. et al. Stable C0T-1 repeat RNA is abundant and is associated with euchromatic interphase chromosomes. Cell 156, 907–19 (2014).

72. Hall, L.L. et al. An ectopic human XIST gene can induce chromosome inactivation in postdifferentiation human HT-1080 cells. Proc Natl Acad Sci U S A 99, 8677–82 (2002).

73. Minks, J. & Brown, C.J. Getting to the center of X-chromosome inactivation: the role of transgenes. Biochem Cell Biol 87, 759–66 (2009).

74. Namekawa, S.H., Payer, B., Huynh, K.D., Jaenisch, R. & Lee, J.T. Two-step imprinted X inactivation: repeat versus genic silencing in the mouse. Mol Cell Biol 30, 3187–205 (2010).

75. Platt, E.J., Smith, L. & Thayer, M.J. L1 retrotransposon antisense RNA within ASAR lncRNA genes controls chromosome-wide replication timing. Journal of Cell Biology in press(2017).

76. Figueroa, D.M., Darrow, E.M. & Chadwick, B.P. Two novel DXZ4-associated long noncoding RNAs show developmental changes in expression coincident with heterochromatin formation at the human (Homo sapiens) macrosatellite repeat. Chromosome Res 23, 733–52 (2015).

77. Elisaphenko, E.A. et al. A dual origin of the Xist gene from a protein-coding gene and a set of transposable elements. PLoS One 3, e2521 (2008).

78. Kruskal, J.B. Multidimensional scaling by optimizing goodness of fit to a nonmetric hypothesis. Psychometrika 29, 1–27 (1964).

79. Zhu, Y., Bye, S., Stambrook, P.J. & Tischfield, J.A. Single-base deletion induced by benzo[a]pyrene diol epoxide at the adenine phosphoribosyltransferase locus in human fibrosarcoma cell lines. Mutat Res 321, 73–9. (1994).

80. Dobin, A. et al. STAR: ultrafast universal RNA-seq aligner. Bioinformatics 29, 15–21 (2013).

81. Li, H. et al. The Sequence Alignment/Map format and SAMtools. Bioinformatics 25, 2078–9 (2009).

82. Marchal, C. et al. Genome-wide analysis of replication timing by next-generation sequencing with E/L Repli-seq. Nat Protoc 13, 819–839 (2018).

83. Li, H. & Durbin, R. Fast and accurate short read alignment with Burrows-Wheeler transform. Bioinformatics 25, 1754–60 (2009).

84. Smith, L., Plug, A. & Thayer, M. Delayed Replication Timing Leads to Delayed Mitotic Chromosome Condensation and Chromosomal Instability of Chromosome Translocations. Proc Natl Acad Sci U S A 98, 13300–13305 (2001).

85. Smith, L. & Thayer, M. Chromosome replicating timing combined with fluorescent in situ hybridization. J Vis Exp, e4400 (2012).

